# Rapid diffusion of large TatA complexes detected using single particle tracking microscopy

**DOI:** 10.1101/2020.05.14.095463

**Authors:** Aravindan Varadarajan, Felix Oswald, Holger Lill, Erwin J.G. Peterman, Yves J. M. Bollen

## Abstract

The twin-arginine translocation (Tat) system transports folded proteins across the cytoplasmic membrane of most bacteria and archaea. TatA, which contains a single membrane-spanning helix, is believed to be responsible for the actual translocation. According to the prevalent model, multiple TatA subunits form a transient protein-conducting pore, which disassembles after each translocation event. An alternative model exists, in which TatA proteins locally weaken the lipid bilayer to translocate folded proteins. Here, we imaged eGFP-fused TatA expressed from the genome in live *E. coli* cells. Images showed TatA occuring both in highly mobile monomers or small oligomers and in large, stable complexes that do not dissociate. Single-particle tracking revealed that large TatA complexes switch between fast and slow diffusion. The fast diffusion is too fast for a transmembrane protein complex consisting of multiple TatA monomers. In line with recent data on rhomboid proteases, we propose that TatA complexes switch between a slowly diffusing transmembrane conformation and a rapidly diffusing membrane-disrupting state that enables folded proteins to cross the membrane, in accordance with the membrane-weakening model.

## Introduction

Bacteria and archaea use two protein transport systems for translocating proteins across the cytoplasmic membrane (Natale, Brüser et al., 2008). The universal secretory (Sec) system transports unfolded proteins through a narrow pore formed by the SecYEG protein complex in the membrane while hydrolyzing ATP, whereas the Twin arginine transport (Tat) system transports folded proteins by an uncharacterized mechanism, apparently driven solely by proton motive force (pmf) (Natale et al., 2008). One of the key properties of the Tat system is its ability to translocate fully folded proteins across the cytoplasmic membrane without substantial leakage of other cytoplasmic content into the periplasm.

The Tat pathway is not found in all bacteria and archaea. However, in those that do contain the system, it plays a crucial role in various cellular processes, including energy metabolism, formation of the cell envelope, and nutrient acquisition (Palmer & Berks, 2012). It has been shown that the Tat system is essential for the virulence of several bacterial pathogens (Alcock, Baker et al., 2013, Pradel, Ye et al., 2003). In plant chloroplasts, which are evolutionarily linked to bacteria, the Tat pathway plays an essential role in the biogenesis of the photosynthetic electron-transport chain. Substrate proteins that are translocated by the Tat pathway are synthesized with an amino-terminal extension that has a consensus sequence motif, containing two highly conserved consecutive arginine residues (Alcock et al., 2013). Mutation studies have shown that substrate translocation is impaired when these arginine residues are substituted with other amino acid residues (Frobel, Rose et al., 2012, Stanley, Palmer et al., 2000). In the gram-negative bacterium *Escherichia coli*, the Tat system consists of four integral membrane proteins TatA, TatB, TatC and TatE (De Leeuw, Porcelli et al., 2001). TatA, TatB and TatE are structurally related, they all contain an amino-terminal hydrophobic helix followed by an amphipathic helix and an unstructured domain. The hydrophobic helix of these proteins is too short to span the membrane, a property that has been proposed to play a role in the translocation mechanism (Rodriguez, Rouse et al., 2013), which is still largely unknown. TatB and TatC form large complexes (400 – 600 KDa) containing several copies of both proteins in a 1:1 ratio (Bolhuis, Mathers et al., 2001). These TatBC complexes interact with the N-terminal signal peptide of Tat substrates and act as substrate receptors (Alami, Luke et al., 2003, Cline & Mori, 2001). TatA is the most abundant protein of the Tat system and is thought to be its pore-forming component (Lee, Tullman-Ercek et al., 2006). TatE, a functional homologue of TatA is less abundant. It can partially compensate for the lack of TatA activity in a knock-out strain (Alcock et al., 2013). For long, it was believed that under most laboratory conditions TatE does not play an important role in translocation. Recently, it has, however, been shown that a GFP fusion to TatA, which in absence of TatE results in heavily compromised Tat activity, is much better tolerated in the presence of intact TatE (Alcock et al., 2013, Eimer, Frobel et al., 2015).

Detergent-solubilized TatA forms large homo-oligomeric complexes of variable size (Oates, Barrett et al., 2005, Porcelli, de Leeuw et al., 2002). Cryo-electron microscopy has revealed that TatA complexes form ring-like shapes with an internal cavity wide enough to transport folded substrate proteins. Single-molecule imaging in live *E. coli* cells showed that there are about 100 TatA monomers freely diffusing and 15 TatA complexes present in the membrane (Leake, Greene et al., 2008). These complexes have a wide distribution of sizes, with an average of 25 TatA monomers per complex (Leake et al., 2008). Recently, it has been shown in *E. coli* that increasing the number of Tat-transportable substrate proteins increases the number of TatA complexes per cell. Formation of these TatA complexes requires both a functional TatBC substrate receptor and the transmembrane pmf (Alcock et al., 2013). These observations have led to the “substrate-induced Tat association model” (Alcock et al., 2013). In this model the assembly of an active Tat complex begins with binding of substrate to the TatBC complex, possibly via 2D diffusion on the membrane surface (Shanmugham, Wong Fong Sang et al., 2006), followed by pmf-dependent recruitment of TatA, which is dispersed in the membrane as monomers or small oligomers. Subsequently, TatA forms a pore through which substrate protein can pass from the cytoplasmic to the periplasmic side. Finally, after translocation, the signal peptide is cleaved from the substrate and the Tat complex disassembles, resetting the translocation cycle.

If this model is correct, one would expect that (i) TatA complexes assemble and disassemble during the translocation cycle and (ii) that increase or depletion of Tat substrate alters TatA complex number and size. To test these predictions, we performed a combination of single-molecule fluorescence experiments and biochemical assays on an *E. coli* strain containing TatA-eGFP at the endogenous TatA locus in the genome under various conditions: with natural expression levels of Tat substrate proteins, overexpressing a substrate, overexpressing a transport-arresting substrate, and with inhibited protein synthesis. We find that the average number of TatA complexes per bacterium indeed increases with substrate load, in accordance with the model and with previous findings. However, these complexes are stable and do not dissociate under any condition tested. We furthermore find that the diffusion of TatA complexes is heterogeneous under all conditions: complexes switch between slow and very fast diffusion. The fast diffusion coefficient is substantially larger than would be expected for a large protein complex embedded in the inner membrane. Furthermore, a considerable amount of TatA can be washed off the membrane surface with a high-pH treatment. We discuss how these observations may relate to the structure of TatA complexes and their role in the Tat translocation process.

## Results

### Fluorescently labeled genomic TatA is functional

We constructed an *E. coli* strain in which the native tatA gene within the genome was replaced by a gene encoding a TatA-eGFP fusion. The resulting strain, referred to as TatA^eGFP^BC, lacks the native TatA protein but expresses TatA-eGFP, as well as TatB, TatC, and TatE from the chromosome under control of the endogenous promoter.

We assessed the transport activity of the TatA^eGFP^BC strain by examining the translocation of the Tat substrate protein SufI fused to the HA-tag (Jong, ten Hagen-Jongman et al., 2004). SufI, a 55 kD cell division protein that is involved in the assembly of the bacterial divisome, is transported from cytoplasm to periplasm via the Tat system. During the transport process, the N-terminal signal peptide (~5 kD) of SufI is cleaved off at the outer surface of the inner membrane, resulting in the release of a 50 kD protein in the periplasm. Western blotting of SufI-HA expressed from Tat knockout strains (TatA/E deletion strain Jarv16 as well as a TatABC deletion strain) showed a prominent 55 kD band indicating that SufI was not processed and hence not transported to the periplasm (Fig. 1A). In contrast, the TatA^eGFP^BC and wild-type MC4100 *E. coli* strains showed an intense 50 kD and a less intense 55 kD band indicating that most of the SufI produced in these strains was processed and transported to the periplasm (Fig. 1A).

**Figure 1.**
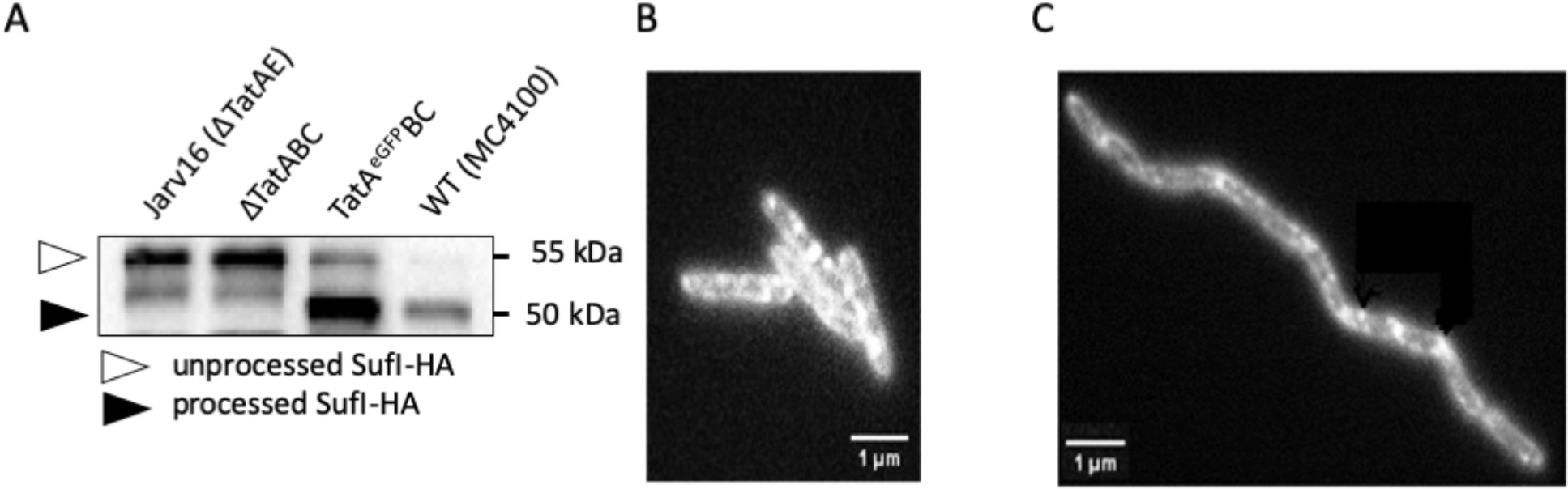
Functionality of TatA^eGFP^. (A) Western blot of whole-cell lysates from MC4100 *E. coli* strains expressing SufI-HA substrate protein. Lysates of Jarv16, ΔTatABC, TatA^eGFP^BC, and wild-type MC4100 are probed with anti-HA antibodies. White arrow indicates unprocessed SufI-HA (55 kD) and black arrow indicates processed SufI-HA (50 kD). B, C. Wide-field fluorescence microscopy image of TatA^eGFP^BC *E. coli* cells. (B) Cells expressing functional TatA-eGFP from the genome. (C) Cells expressing TatA-eGFP in a TatB truncation background. Cells are aspecifically immobilized on a glass slide. High fluorescence intensity at the edges of the cells indicates membrane localization of TatA-eGFP.

TatA^eGFP^BC cells were examined using epi-illuminated wide-field fluorescence microscopy. Bacteria appeared to have fluorescent contours containing bright spots, indicating that TatA-eGFP localizes to the membrane and forms complexes (Fig. 1B). In addition, cells were well separated and had normal shapes, whereas cells without functional TatA are known to suffer from elongated growth and chain formation (Fig. 1C) (Stanley, Findlay et al., 2001). Together, these results shown in Figure 1 indicate that the function of TatA is not hampered by eGFP fusion, which is in agreement with previous observations (Alcock et al., 2013, Frobel et al., 2012, Rose, Frobel et al., 2013).

### TatA complex formation is substrate dependent whereas disassembly is not

Fluorescence images of cells expressing TatA-eGFP (Fig. 1B) show large complexes as well as a diffuse background, which is probably due to monomers or small oligomers of TatA-eGFP (Deich, Judd et al., 2004, Leake et al., 2008, Oswald, Bank et al., 2014, van den Wildenberg, Bollen et al., 2011). It has been shown before that an increase of the available Tat substrates results in an increased number of TatA complexes in the membrane (Alcock et al., 2013, Rose et al., 2013). To verify this observation in our strain, we counted the number of TatA complexes in TatA^eGFP^BC cells overexpressing substrate protein SufI from a L-arabinose inducible plasmid. The number of TatA complexes increased gradually from 0 to 2 per cell (average of 1.2) with only naturally expressed substrates to 3 to 6 (average of 3.6) per cell in cells expressing saturating amounts of SufI (Fig. 2A, Supplementary figure 1), indicating that in the presence of more substrate proteins the number of large TatA complexes is increased.

**Figure 2.**
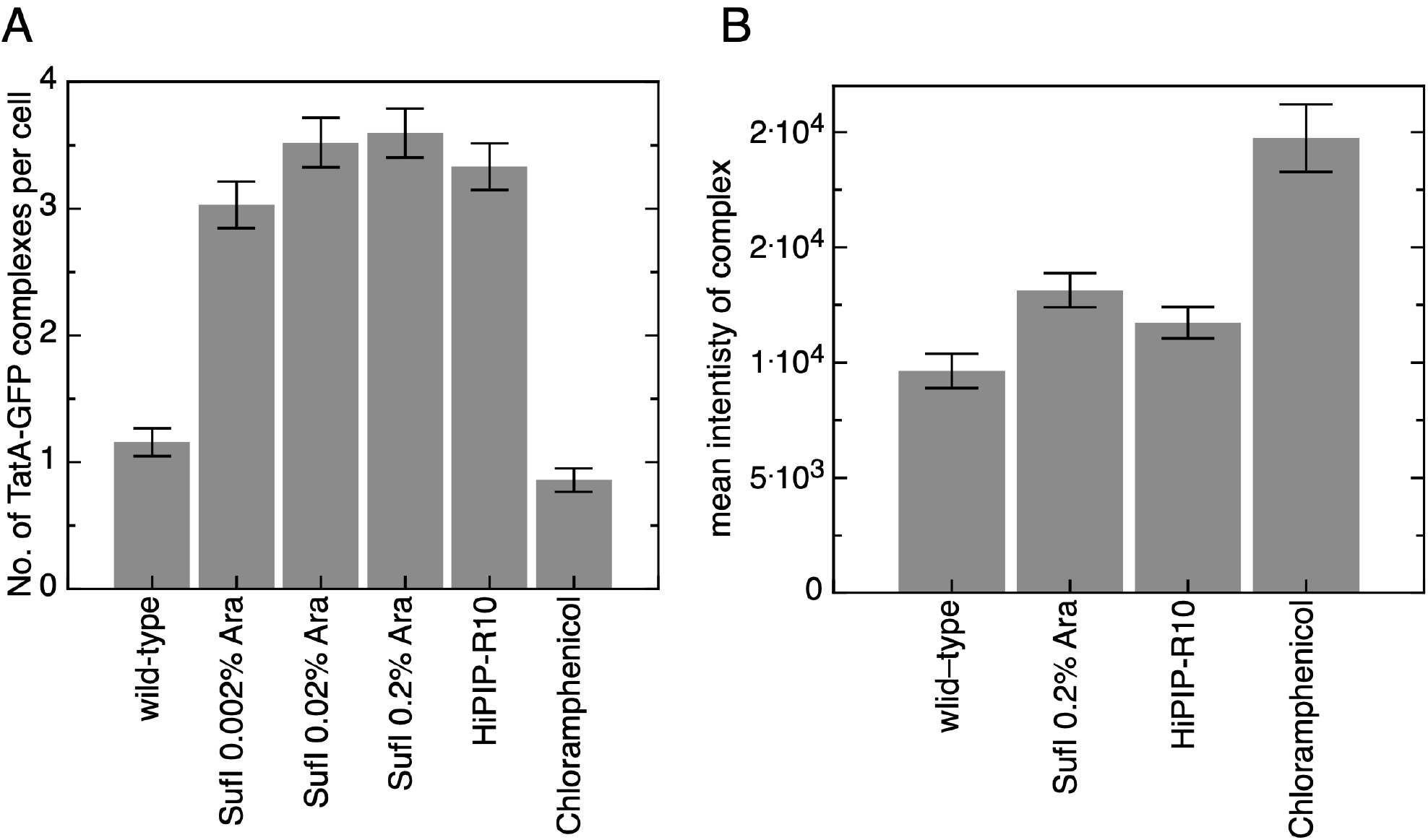
Quantification of TatA-eGFP complexes. (A) The number of TatA-eGFP complexes formed in MC4100 *E. Coli* strain TatA^eGFP^BC expressing wild-type levels of TatA substrates, overexpressing substrate protein SufI at various induction levels, overexpressing transport arresting substrate protein HiPIP-R10 (induced with 0.2% rhamnose), and treated with 100 μg/ml chloramphenicol are represented in the bar graph. Results represent the means of 90 to 100 cells with standard error of the mean (SEM) estimated using 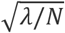, with *λ* the mean and *N* the sample size. Histograms showing the distrubution of counts are shown in supplementary figure 1. (B) Bar graph showing the fluorescence intensities of tracked TatA-eGFP complexes *in vivo* under various conditions, with the stadard error of the mean indicated. For comparison, the mean intensity of purified eGFP under identical imaging conditions is 510 ± 150 a.u. (*N*=183). Histograms showing the distribution of complex intensities are presented in supplementary figure 2.

We also determined the number of TatA-eGFP complexes in bacteria overexpressing a transport-arresting substrate, consisting of high-potential iron-sulfur protein (HiPIP) with a long flexible linker between the signal peptide and the mature domain (Lindenstrauss & Brüser, 2009, Taubert & Brüser, 2014). These HiPIP-R10 overexpressing cells contained on average more than 3 large, bright TatA complexes, similar to cells overexpressing SufI at a comparable level of induction (Fig. 2A, Supplementary figure 1).

An increase in the number of TatA-eGFP complexes might be caused by either an increase in TatA expression level, or by a re-distribution of the TatA molecules, for example by a shift from monomers and small oligomers to larger complexes or by decreasing the average size of a complex. In order to discriminate between these two possibilities, we measured the expression level of TatA-GFP by using flow cytometry under wild-type condition, in cells overexpressing SufI, and in cells overexpressing HiPIP-R10. The three distributions of total cellular fluorescence intensity were nearly indistinguishable (Supplementary figure 3). Hence, these results indicate that the increase in the number of TatA complexes results from a re-distribution of a relatively constant pool of TatA monomers.

It has been hypothesized that TatA complexes disassemble after every translocation cycle (Alcock et al., 2013). To test this hypothesis, we treated TatA^eGFP^BC wild-type strain with chloramphenicol for ~3 hours. Chloramphenicol blocks protein synthesis on the ribosome, which presumably results in complete translocation of the remaining, already synthesized Tat substrate proteins, and eventually in depletion of substrate protein in the cytoplasm. We imaged these cells and determined the average number of TatA complexes. Strikingly, the chloramphenicol treated cells contained on average one large TatA complex per cell, only slightly less than the wild-type condition, suggesting that TatA complexes do not disassemble in the absence of substrate proteins, at least not within several hours.

### TatA complex size varies with substrate condition

The TatA complexes in cells treated with chloramphenicol appeared to be much brighter than the TatA complexes under other conditions. In order to obtain more quantitative insight into the size distributions of the TatA-eGFP complexes under various substrate conditions, we quantified the fluorescence intensities of individual TatA-eGFP complexes. Under wild-type substrate conditions the intensity of TatA-eGFP complexes shows a rather broad distribution with a mean intensity that corresponds to 19 TatA-eGFP molecules per complex. Upon over-expression of a substrate - either the transport-competent SufI or the transport-arresting HiPIP-R10, the intensity of the complexes increases, with mean intensities corresponding to 26 and 23 TatA-eGFPs per complex, respectively (Fig. 2B, supplementary figure 2). These results are consistent with an electron microscopy study of purified TatA complexes, which showed that TatA forms complexes of various sizes, including large ones in the range of 20-35 monomers (10-13 nm diameter) (Gohlke, Pullan et al., 2005). Chloramphenicol-treated cells contained substantially larger complexes containing 39 TatA-eGFPs on average (Fig. 2B).

### The diffusion of TatA complexes has an unexpected size dependence

TatA-eGFP fluorescence intensity reports on the number of TatA proteins in a complex. However, TatA is known to also interact with TatB and TatC in a substrate-dependent manner. Since in general, the diffusion coefficient of a trans-membrane protein complex depends on its size (Oswald, Varadarajan et al., 2016), we reasoned that diffusion analysis of TatA-eGFP complexes might reveal those interactions with unlabeled membrane proteins. To this end, we tracked individual TatA-eGFP complexes under wild-type Tat-substrate conditions, in cells overexpressing the transportable substrate SufI and in cells overexpressing the transport-arresting substrate HiPIP-R10. Using automated single-particle tracking, stacks of images were analyzed and diffusion trajectories of large, bright TatA complexes were reconstructed. Figure 3 shows an example of two cells in which bright mobile fluorescent spots are tracked (panel A) and of one reconstructed trajactory (panel B). These data are also available as a movie (supplementary movie 1).

**Figure 3.**
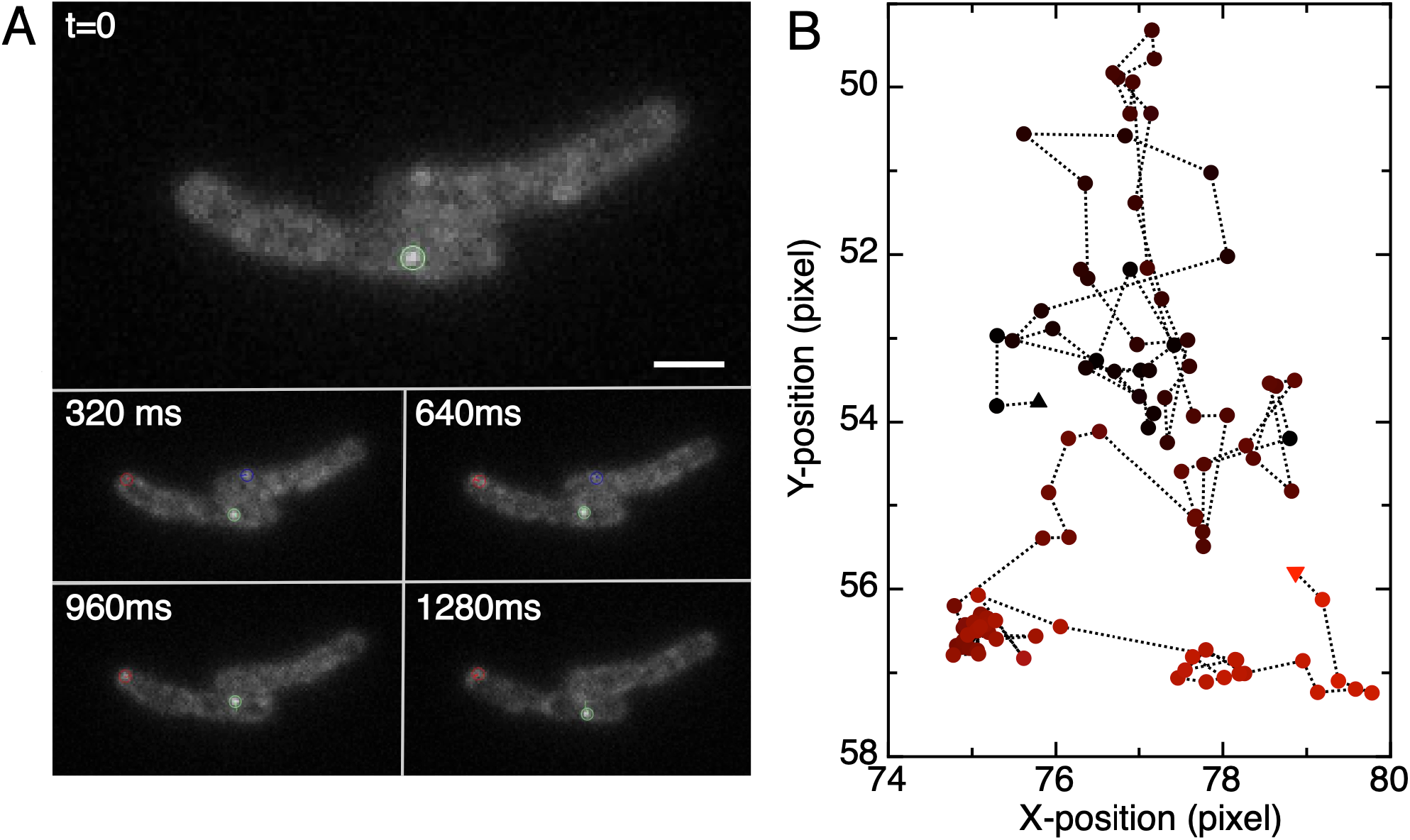
Tracking of diffusing TatA-GFP complexes. Fluorescence images are recorded with 32 ms exposure. (A) Fluorescence images are shown at various time intervals. Detected spots are indicated with colored circles. The scale bar is 1 μm. (B) Reconstructed trajectory of the particle shown in green in panel A. Time is indicated by color, from black to red. First and last localisation are shown by triangles. Position is indicated as pixel; one pixel in the image corresponds to 80 nm.

The trajectories obtained from microscopy are 2D projections of the actual 3D trajectories along the highly curved membrane surface. Therefore, direct analysis of this 2D-projected data would yield erroneous results (Deich et al., 2004, Leake et al., 2008, Oswald et al., 2014, van den Wildenberg et al., 2011). To avoid this, we applied IPODD (Inverse Projection Of Displacement Distributions) (Oswald et al., 2014) to reconstruct 3D displacements and determine diffusion coefficients using mean squared displacement (MSD) analysis. We determined that the diffusion coefficients of TatA-eGFP complexes in wild-type substrate conditions (*D* = 0.099 ± 0.002 μm^2^ s^−1^) and SufI substrate overexpressing conditions (*D* = 0.094 ± 0.003 μm^2^ s^−1^) were nearly identical (Fig. 4A, Table 1). In contrast, when overexpressing the transport-arresting substrate HiPIP-R10, the diffusion coefficient of TatA-eGFP complexes was 2.5-fold lower (*D* = 0.039 ± 0.001 μm^2^ s^−1^). Furthermore, after chloramphenicol treatment, TatA diffusion instead increased 1.6-fold compared to wild-type substrate conditions (*D* = 0.156 ± 0.003 μm^2^ s^−1^). A plot of the diffusion coefficient as obtained by MSD analysis against the average number of TatA-eGFP molecules per complex estimated from fluorescence intensity shows a rather unexpected correlation between diffusion coefficient and number of TatA per complex under these four conditions: the conditions with a larger number of TatA per complex, show larger diffusion coefficient (Figure 4B). Typically, the opposite is expected and observed (Oswald et al., 2016, Saffman & Delbruck, 1975): the larger the transmembrane protein complex, the slower the diffusion coefficient.

**Table 1.** Diffusion coefficients (*D*) with relative occurrence (γ) that best describes the diffusive behavior of TatA-eGFP under various conditions. Values are average ± standard deviation obtained from two independent experiments.

| Substrate condition | $D_{MSD} (\mu m^2/s)$ | $D_1 (\mu m^2/s)$ | $D_2 (\mu m^2/s)$ | $\gamma(D_1)$ |
| --- | --- | --- | --- | --- |
| wild-type | $0.099 \pm 0.002$ | $0.024 \pm 0.005$ | $0.175 \pm 0.005$ | $0.57 \pm 0.02$ |
| Over-expression of SufI<br>(induced with 0.2% Ara) | $0.094 \pm 0.003$ | $0.023 \pm 0.002$ | $0.184 \pm 0.003$ | $0.61 \pm 0.03$ |
| Over-expression of HiPIP<br>(induced with 0.2% Rham) | $0.039 \pm 0.001$ | $0.024 \pm 0.003$ | $0.117 \pm 0.004$ | $0.82 \pm 0.02$ |
| TatA<sup>eGFP</sup>BC treat with<br>Chloramphenicol (100 $\mu g/ml$) | $0.156 \pm 0.003$ | $0.049 \pm 0.008$ | $0.230 \pm 0.015$ | $0.55 \pm 0.05$ |

**Figure 4.**
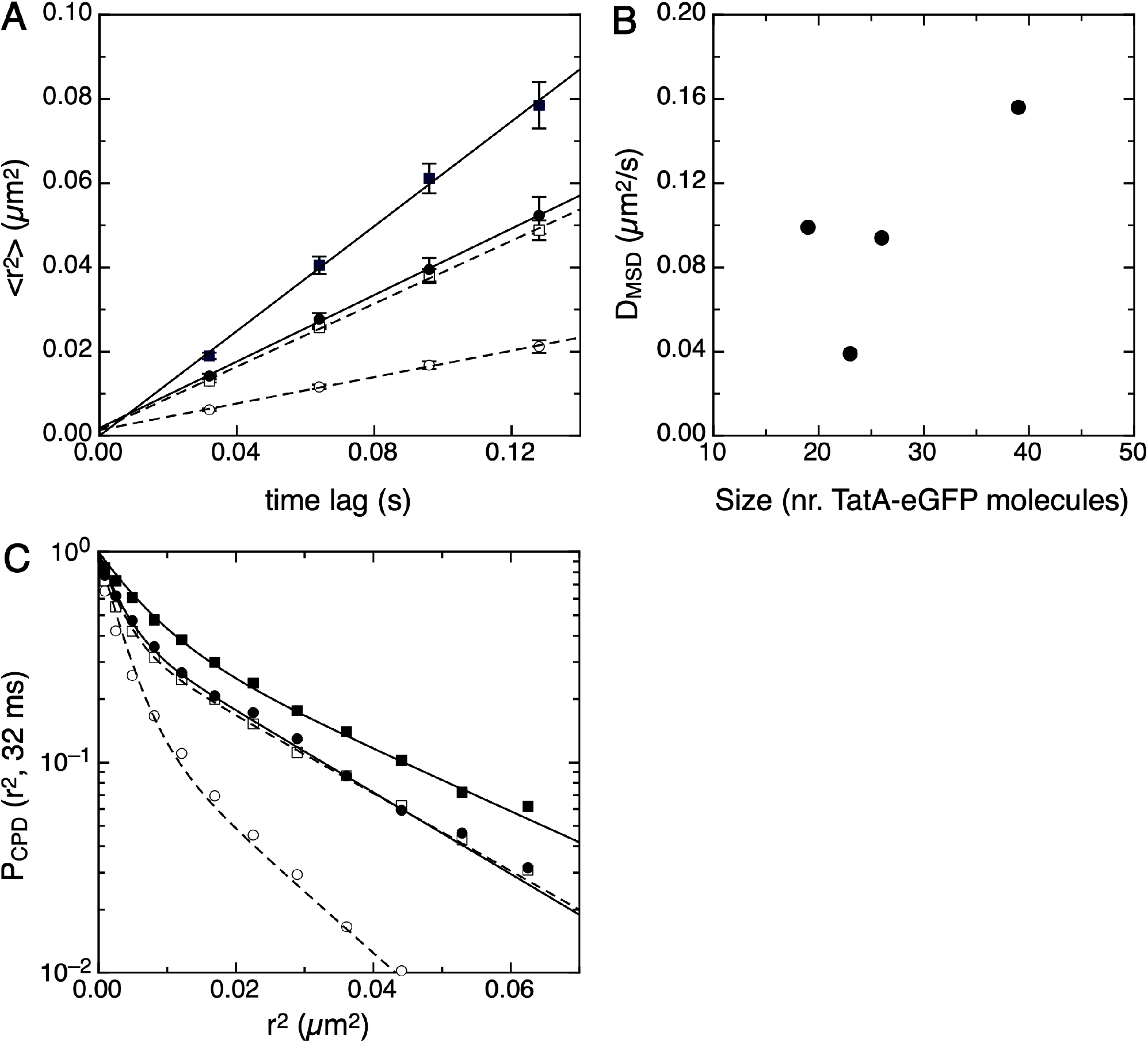
Diffusion analysis of TatA-eGFP complexes in the membrane of living bacteria. (A) Mean squared displacement plotted against time lag for Wild-type substrate conditions (filled circles), substrate protein SufI overexpression (open squares), transport arresting substrate protein HiPIP-R10 overexpression (open circles) and chloramphenicol treatment (filled squares). Lines: linear fits yielding diffusion coefficients (*D*) (see Table 1). Solid lines correspond to filled symbols, dashed lines correspond to open symbols. (B) A plot of the diffusion coefficients from panel (A) against the average number of TatA molecules per complex from Figure 2B shows that the conditions with more TatA molecules per complex tend to have higher diffusion coefficients. (C) CPD analysis of TatA-eGFP diffusion. Symbols as in panel A. Lines: double exponential fits yielding two populations with distinct diffusion coefficients (see Table 1).

### Diffusion of TatA is heterogeneous

Visual inspection of the trajectory shown in Fig. 3B gives the impression that diffusion of TatA-eGFP commplexes is heterogeneous. The first half of the trajectory, shown in darker colors, looks like normal diffsion. In the second half however, shown in redder colors, the TatA complex appears to swich between normal and very slow diffusion. Further inspection of more image sequences confirmed this impression. It would be interesting to confirm this impression in an objective manner. MSD analysis, being an averaging technique, is not able to reveal heterogeneity. Cumulative probability distribution (CPD) analysis (Sonnleitner, Schutz et al., 1999) in contrast, can unravel heterogeneity. Using this analysis method on the IPODD-corrected displacement distributions, we found that the CPD of TatA-eGFP complex displacements was not following a single exponential decay as is expected for homogeneous diffusion, in all four conditions (Fig. 4C).Instead, the CPDs could be well fitted with a sum of two exponentials (Fig. 4C and Table 1), reflecting the presence of two populations with distinct diffusion coefficients (*D_1_* and *D_2_*), differing an order of magnitude, with relative occurrences *γ* and 1-*γ*, respectively. The TatA-eGFP diffusion parameters under wild-type substrate conditions and overexpressing SufI were almost identical. Overexpressing transport-arresting substrate HiPIP-R10, in contrast, resulted in only slightly different diffusion coefficients, but a substantially larger fraction of slow-diffusing complexes. Chloramphenicol treatment resulted in increases of both diffusion coefficients (2-fold for the slow component, 1.3-fold for the fast), leaving the ratio relatively unaffected. Together these results show that the diffusion of TatA is heterogeneous, in the sense that in all four conditions tested, fast and slow-moving TatA complexes co-exist. Furthermore, we often observed individual particles switching between fast and slow diffusion modes without changing in fluorescence intensity (see for example Fig. 3B and Supplementary movie 1). Switching is observed under all conditions, even without substrate proteins available (see Supplementary movie 2). It is thus not an effect of transport activity, but rather appears to be an intrinsic property of TatA complexes.

We next wondered how the fast and slow diffusion modes correlate with complex size. In other words: which component is responsible for the increase of the average diffusion coefficient with complex size as shown in Figure 4B. However, a clear correlation could not be observed. Both diffusion coefficients appear to increase with the number of TatA-eGFP molecules in a complex, and also the fraction of the fast component increases accordingly: TatA complexes in the presence of chloramphenicol, which are the largest (43 TatA monomers on average), are fast for 45% of the time, and they also have the largest values for D_1_ and D_2_.

### Does TatA occur as a peripheral membrane protein?

In a previous study, we have related transmembrane protein size to diffusion coefficient in live *E. coli* (Oswald et al., 2016). A comparison with that study indicates that the slow TatA diffusion coefficient D_1_ (0.023 - 0.05 μm^2^s^−1^) corresponds to protein complexes of some 20-40 transmembrane helices, consistent with the intensities of the fluorescent spots (Figure 2) and existing data on TatA complex size (Gohlke et al., 2005). The fast diffusion coefficient D_2_ (0.12 - 0.18 μm^2^s^−1^), however, is not in line with TatA complex size. It is even faster than that of WALP-KcsA-eGFP, which contains only two trans-membrane helices and has an ~6.5 times smaller radius than the TatA complexes (Oswald et al., 2016). In other words, the fast diffusion coefficient appears to be inconsistent with such large complex sizes diffusing in the plane of the membrane. Is it possible that large TatA-containing complexes exist in a state that is not embedded in the membrane? *E. coli* TatA is generally considered to be a genuine trans-membrane protein (De Leeuw et al., 2001). Recently however, *E. coli* TatA fused to a Strep-tag was shown to be partially extracted from the inner membrane (Hou, Heidrich et al., 2018, Taubert, Hou et al., 2015). In order to find out if rapid diffusion of large TatA complexes can be explained by them not being properly incorporated in the membrane, we isolated *E. coli* membranes and washed them with sodium carbonate. Sodium carbonate can remove membrane proteins that are not firmly bound to the membrane surface, leaving other, more tightly integrated membrane proteins in place (Molloy, 2008).

Western-blot data revealed that upon washing membranes with sodium carbonate, both fractions (supernatant and membrane pellet) contain a considerable amount of TatA-eGFP (Fig. 5). In contrast, washing the membrane with PBS did not result in detectable amounts of TatA-eGFP in the supernatant fraction. The integral membrane protein MscS was used as control for membrane integrity during carbonate washes. As expected, MscS is not extracted from the membrane by sodium carbonate. Also, the combined cytoplasmic and periplasmic fractions (i.e. the supernatant fraction during the initial membrane isolation) do not contain substantial amounts of TatA or MscS, respectively. Together, these results indicate that TatA-eGFP may be present in two fractions in the cell: one tightly bound to the membrane, and one less tightly bound, but still associated with the membrane. Strikingly, the TatA-eGFP in the supernatant of the carbonate wash experiments was not detected by the anti-TatA antibody (supplementary Figure 4). To rule out that the band in Figure 4 is an artifact of the anti-GFP antibody, the experiment was repeated with a TatA-mCherry strain and an anti-mCherry antibody, which produced similar results as shown in Figure 5 (see supplementary figure 5). Finally, the supernatant fraction of the carbonate washing experiment of TatA-eGFP containing membranes was analyzed using mass spectrometry. Three peptides were found that are unique for eGFP. The sequences of those peptides were FEGDTLVNR, FSVGEGEGDATYGK, and SAMPEGYVQER. The same carbonate washing experiment was performed on wild-type *E. coli* strain MG1655. Western blot analysis with an anti-TatA antibody showed the presence of TatA only in the membrane fraction (supplementary Figure 4), consistent with our observation that the TatA antibody does not recognize TatA outside the membrane. However, mass spectrometry revealed the presence of one unique TatA peptide in the supernatant of the carbonate washing experiment, with the sequence AMSDDEPKQDKTSQADFTA, corresponding to the end of the amphipathic helix and the beginning of the polar tail of TatA. We conclude that a substantial fraction of TatA molecules occurs in a conformation or in a specific membrane environment in which it can be extracted by carbonate, and that is not recognized by antibodies against TatA.

**Figure 5.**
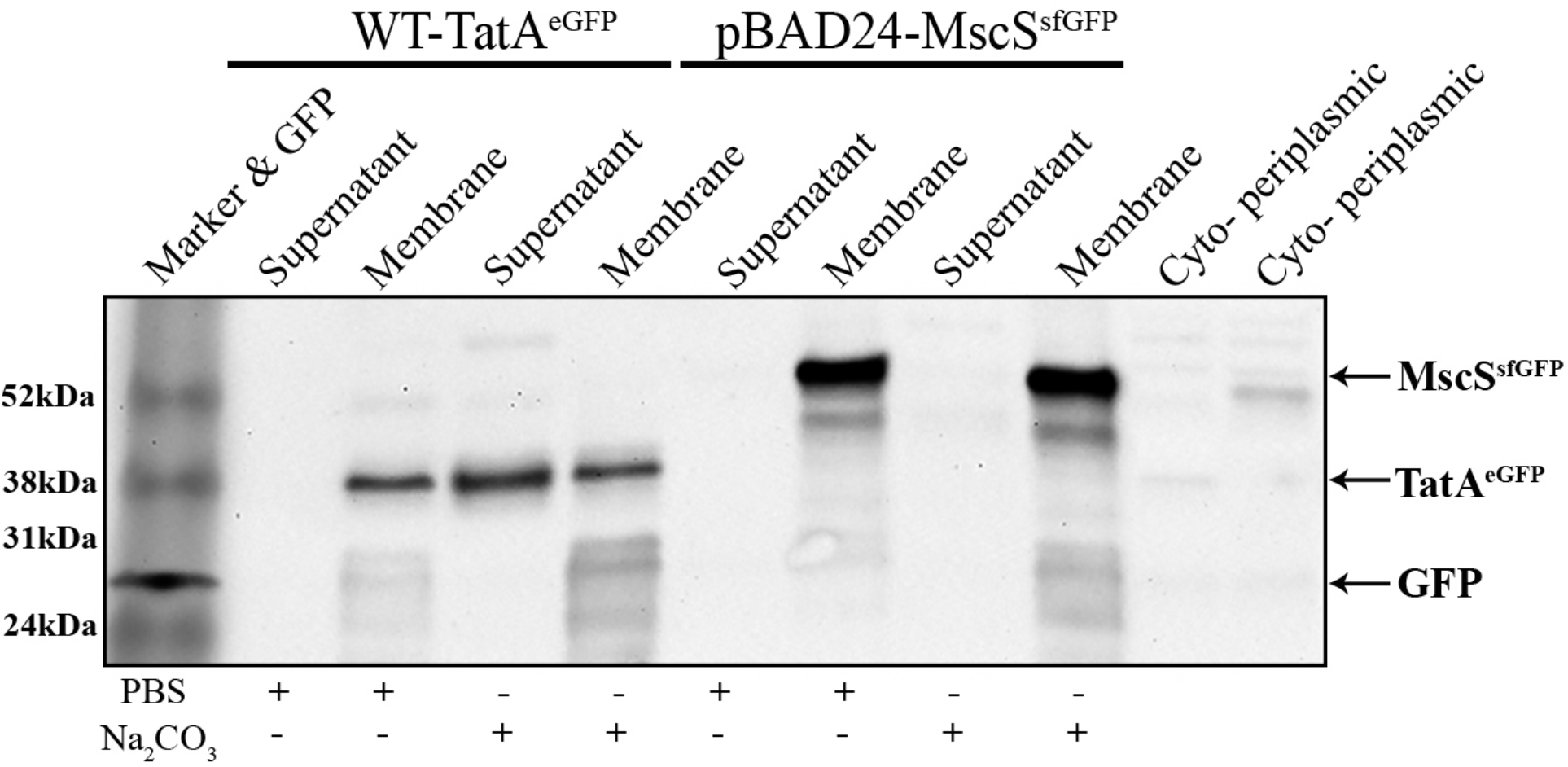
*E. coli* membrane washing detected with western blotting. Membranes were isolated from *E.coli* cells genomically expressing TatA-eGFP and from cells expressing MscS-sfGFP from a plasmid. These membranes were washed with PBS or carbonate buffers as indicated in the bottom, and then analysed using SDS-PAGE followed by western-blotting using an anti-GFP antibody. Purified eGFP(his-tag) was used as a GFP molecular mass indicator as well as an antibody specificity indication.

**Figure 6.**
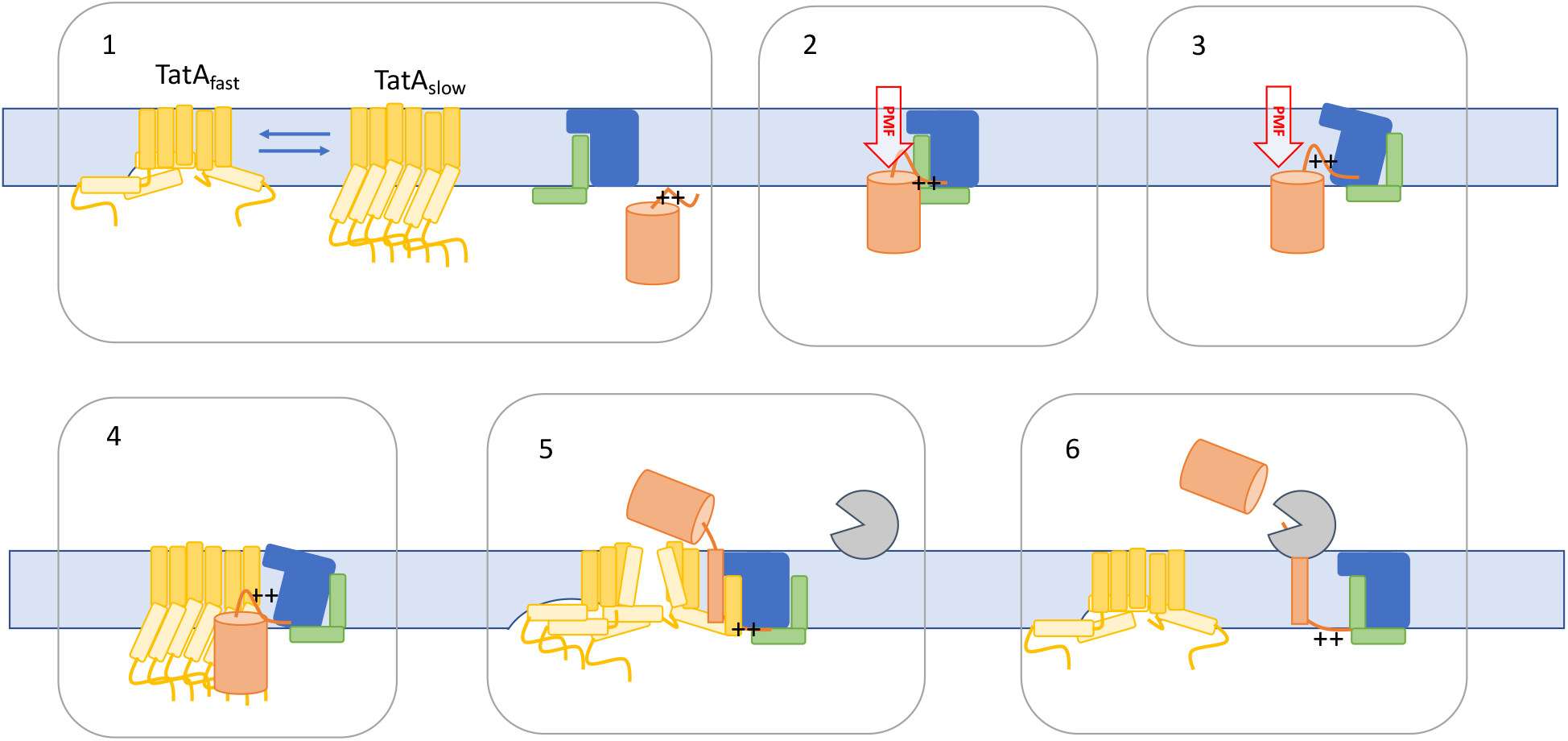
A novel mechanistic model for the Twin Arginine Translocation system. 1. Resting state of the Tat system. TatA consisting of a short trans-membrane helix (dark yellow) and an amphipathic helix (lighter yellow) forms oligomeric complexes that switch between two conformations: a rapidly diffusing one that deforms the membrane, and a slowly diffusing one in which part of the amphipathic helix is pulled into the membrane. TatB (green) and TatC (blue) form a separate receptor complex. For simplicity, TatB and TatC are shown as monomers, although they probably form an oligmeric assembly. A substrate molecule (orange) approaches the TatBC complex, presumably via lateral diffusion over the membrane surface. The two arginine residues are represented by two + signs. 2. The substrate molecule binds to its binding site on the cytoplasmic surface of TatC. The pmf facilitates deeper insertion of the signal peptide into the membrane (this has been shown for thylakoid TatC (Gerard & Cline, 2007)). The two arginines may be pulled into the membrane, leading to a minor deformation of the bilayer. 3. Substrate binding results in a conformational change of TatC, shown here by tilting, causing TatB to move away from the TatA binding site at the short trans-membrane helices 5 and 6 of TatC (Alcock et al., 2016). 4. The TatA complex approach the TatBC complex by diffusion. They may be attracted by the membrane deformation by the short TatC helixes 5 and 6 together with the substrate. Considering that over-expression of HiPIP-R10, which assembles the entire translocation complex but cannot be translocated, leads to much slower diffusion, we propose that it is the slow TatA complex that binds to TatBC initially. 5. Together, TatA and the signal peptide weaken the membrane to such an extent that it ruptures, thus allowing the folded substrate to reach the periplasm, and the signal peptide to form a trans-membrane helix. 6. Leader peptidase (grey) cleaves the signal peptide and releases the folded substrate into the periplasm. TatA dissociates from TatBC, thus allowing TatB to take its original position. The Tat system is back in its resting state, with a TatA complex switching between inserted and associated states. Once assembled, TatA complexes do not dissociate as long as the pmf is not disrupted. TatA complexes keep switching between a rapidly diffusing membrane-disrupting state and a normally diffusing membranespanning state.

## Discussion

TatA appears to be the key component of the Twin Arginine Translocation system, responsible for the actual translocation of folded proteins across the inner membrane. It is believed to form complexes that vary in size during the translocation cycle by first assembling to form a pore complex upon substrate binding to the TatBC oligomer, and then disassemble after translocation is completed (Alcock et al., 2013). Disassembly of the pore complex is believed to aid in preventing accidental opening of the large pore in the absence of a substrate, which could result in leakage of cellular components into the periplasm. Here we used single-molecule fluorescence microscopy to study the size and mobility of individual TatA complexes in living *E. coli* bacteria. Under native substrate conditions, most cells contained one or two large, bright TatA complexes in a diffuse background of monomers or small oligomers. Upon overexpression of (functional) Tat substrate SufI we observed that bacteria contained more TatA-eGFP complexes, while the total fluorescence intensity of the cells did not change. Furthermore, complexes had similar fluorescence intensity and diffusion properties as in strains with wild-type Tat-substrate expression. Together these observations indicate that substrate overexpression does not elevate TatA expression, but rather stimulates the formation of large TatA complexes from a pool of TatA molecules, dispersed in much smaller units within the membrane. Also overexpressing HiPIP-R10, an artificial Tat substrate that results in translocation arrest (Lindenstrauss & Brüser, 2009), leads to an increased number of TatA-eGFP complexes. The fluorescence intensity, however, of these complexes was significantly lower, implying that they contained less TatA. At the same time, these complexes diffused more slowly, indicative for a larger complex diameter. It thus appears that under these conditions Tat complexes are trapped in an intermediate translocation state, containing less TatA molecules and more other membrane proteins, most likely TatB, TatC and the substrate protein.

The observed substrate-dependent complex formation concurs with the substrate-induced association model for the Tat system. However, alternating assembly and disassembly of TatA complexes was not observed here at the experimental time scale of several seconds. Even more, bacteria treated with chloramphenicol contained on average one TatA-eGFP complex, which was substantially larger than under wild-type substrate conditions. This implies that complexes, once formed, are stable for many hours, also after substrate depletion. This observation is not in agreement with the substrate-induced association model. In previous studies, dissociation of TatA complexes is only observed upon depolarization of the membrane by the addition of a chemical uncoupler (Alcock et al., 2013, Rose et al., 2013). Although initial TatA complex formation does seem to be substrate dependent, dissociation into monomers or small oligomers does not appear to occur after each translocation event.

Tracking of TatA-GFP complexes allowed us to quantify their diffusion under various substrate conditions. Mean squared displacement analysis revealed an unexpected correlation between diffusion and complex size: the larger the average complex size in an experiment was, the higher the mean diffusion coefficient. A more detailed analysis of the diffusion using cumulative probability distributions revealed heterogeneous diffusion behavior under all Tat substrate conditions: fast and slow diffusion of large TatA complexes were observed simultaneously, and frequently complexes were seen to switch between fast and slow diffusion. Under all four conditions tested, large TatA-eGFP complexes consisting of up to 50 monomers are seen to diffuse with a diffusion coefficient of around 0.2 μm^2^/s (Table 1). This implies a conceptual discrepancy, since such a fast diffusion coefficient would theoretically correspond to a complex diameter of around 1 nm (Oswald et al., 2016), which can only accommodate one or two trans-membrane helices.

What causes large TatA complexes to diffuse so rapidly? This question triggered us to have a closer look at the presumed trans-membrane orientation of TatA. *E. coli* TatA is generally considered to be a genuine trans-membrane protein (De Leeuw et al., 2001). Some more recent data however, showed that *E. coli* TatA fused to a Strep-tag can be partially extracted from the inner membrane, and that the extractable fraction depends on the growth phase of the cells (Hou et al., 2018, Taubert et al., 2015). Also in other bacteria, such as *Bacillus subtilis* (Pop, Westermann et al., 2003) and *Streptomyces lividans* (De Keersmaeker, Van Mellaert et al., 2005a, De Keersmaeker, Van Mellaert et al., 2005b), a substantial portion of TatA is found outside the membrane. Our Western-blot and mass spectrometric analysis of sodium-carbonate washed membrane revealed that a substantial fraction of TatA-eGFP is not firmly embedded in the cytoplasmic membrane. We did find evidence by mass spectrometry that TatA without eGFP fused to its carboxy terminus can also be extracted from the membrane by carbonate buffer. However, this could not be quantified because the membrane-extracted form of TatA is not recognized by the available antibodies against TatA. It has been previously reported that proteins that are anchored to the membrane by a single membrane-spanning helix can be (partially) extracted from the membrane by carbonate buffer if the transmembrane helix is not very hydrophobic, and if there is no folded domain at the exterior that helps anchoring the protein firmly in the membrane (Kim, Botelho et al., 2015). Considering that TatA has a short transmembrane helix and not a single polar amino acid that protrudes from the membrane at the periplasmic side, it is not such a surprise that some TatA molecules can be extracted by carbonate buffer, and that fusion of a GFP or Strep-tag may influence this delicate balance.

The classical interpretation of carbonate extraction is that those proteins are not properly integrated in the membrane, but are only associated with it, for example via interactions with other proteins or with lipid head groups (Molloy, 2008). For TatA, this would imply that a part of the complexes are not in a trans-membrane orientation. If these membrane-extractable TatA complexes would correspond to the rapidly diffusing fraction of TatA, they would also need to re-enter the membrane, since fluorescence microscopy showed that complexes switch between rapid and slow diffusion. All in all, TatA complexes leaving the membrane as a regular part of the translocation cycle is not a very likely explanation for the rapid diffusion and carbonate sensitivity that we observed.

We therefore saught for an alternative explanation for the rapid diffusion of TatA complexes, assuming that TatA complexes remain properly embedded in the membrane. Recently, Kreutzberger et al. (Kreutzberger, Ji et al., 2019) showed that rhomboid proteases diffuse extremely rapidly through biological membranes, at much higher rates than expected considering their size. It is the mismatch between the short hydrophobic helices of rhomboid proteases and the thickness of the membrane that locally destabilizes the membrane and thereby leads to the extremely rapid diffusion. The same principle may well explain the rapid diffusion that we reported here for large TatA complexes, as well as the fact that part of the complexes are more sensitive to carbonate extraction. Structural analysis and molecular dynamics simulation have shown that the amino-terminal hydrophobic helix of TatA is too short to adopt a stable trans-membrane configuration (Rodriguez et al., 2013). Rodriguez et al. proposed an equilibrium between two conformations for TatA: one in which the TatA proteins span the entire membrane by partial insertion of the amphipathic helix into the membrane, and one in which the amphipathic helix lies on the surface of a thinner, deformed membrane, where larger TatA complexes shift the balance towards membrane deformation (Rodriguez et al., 2013). Here, we frequently observed large TatA complexes switching between rapid and slow diffusion in both directions, and larger complexes have a higher mean diffusion constant. These results agree with the proposed conformations of Rodriguez: slowly diffusing TatA complexes could well correspond to the membrane-inserted state with deformed amphipathic helices and a normal membrane, wheras the rapidly diffusing complexes could correspond to the membrane-deforming conformation of TatA. This conformational switching of TatA also explains the observation that part of the TatA molecules are extracted by carbonate: TatA proteins that are in a locally weakened, thinner membrane are more prone to be extracted than TatA proteins that reside in a proper trans-membrane configuration.

Membrane deformation and the resulting rapid diffusion of TatA appears to be functionally relevant. Already in 2003, membrane deformation has been proposed a possible translocation mechanism of the twin arginine translocation system (Brüser & Sanders, 2003, Rodriguez et al., 2013). Interestingly, rapid diffusion is even observed in cells exposed to chloramphenicol for a long time, and is thus not dependent on binding or translocation of a substrate protein. Rather, it appears to be an intrinsic property of TatA. Large TatA complexes, containing many too short hydrophobic helices, are apparently so unstable in the membrane, that they switch between two energetically nearly equivalent states. The largest complexes that we observed - following substrate depletion by treatment with chloramphenicol - showed the highest average diffusion coefficient. The larger the TatA complexes are, the larger their potential to deform the membrane and switch to the rapidly diffusing state.

How exactly do the observations described here link to the translocation function of TatA? First of all, substrate-induced association of TatA from monomers or small oligomers into larger complexes does occur initially, as observed here and in previous studies by the increase of the number of complexes per cell upon over-expression of a substrate. However, this does not appear to be part of the translocation cycle since dissociation of TatA complexes is not observed. Once formed, TatA complexes are stable and switch between a highly mobile membrane-deforming conformation and a slowly diffusing conformation with the amphipathic helices partly embedded in the membrane. These TatA complexes may associate with a TatBC complex, probably in a susbstrate-dependent manner. These complexes then use the membrane-deforming power of TatA to allow translocation of substrate proteins through the weakened membrane. After translocation, the complex dissociates into a TatBC complex and a TatA oligomer that keeps switching beween rapid and slow diffusion. A visualization of how the Tat system might function based on the above is shown in Fig. 5. The model is adapted from (Alcock, Stansfeld et al., 2016).

## Materials and Methods

### Generation of a ΔTatABC strain

Because correct expression levels of the Tat system have been shown to be crucial for Tat activity (Xiong, Santini et al., 2007), we introduced an in-frame eGFP fusion behind TatA while keeping the Tat-operon structure intact. We followed the two-step homologous recombination protocol from the Court lab (Sharan, Thomason et al., 2009, Thomason, Court et al., 2007). In the first step, the *tatABC* operon was replaced by a double selection marker; in the second step the marker was replaced by *tatA(gfp)BC*. For recombination, we made use of the elements (Exo, Beta and Gam enzymes) of the defective λ prophage, available under a temperature-inducible promoter from the pSIM6 plasmid, which also carries an ampicillin resistance gene (Datta, Costantino et al., 2006, Yu, Ellis et al., 2000). pSIM6 was transformed into *E. coli* strain MC4100. Cells were grown to mid-log phase at 32 °C, after which expression of the homologous recombination genes were induced at 42 °C for 15 minutes. Cells were then cooled on ice, made competent, and transformed with a doublestranded DNA fragment encoding a chloramphenicol acetyl transferase / levansucrase (CAT/SacB) gene cassette, flanked on either side by approximately 40 base pairs of the up- and downstream region of the Tat operon. The CAT/SacB cassette allows for positive selection on Chloramphenicol and counter-selection on sucrose. To check whether the recombination of CAT/SacB cassette was successful, the cells were grown and plated on double-selective Lysogeny Broth (LB) plates containing ampicillin (100 μg/ml) and chloramphenicol (15 μg/ml). The colonies that grew on double selective plates were then cultured and plated on 5% sucrose plates. Finally, colonies that grew on double selective plates and died on sucrose plate were screened using Polymerase Chain Reaction (PCR) to identify positive ΔTatABC colonies.

### Recombination of TatA^eGFP^BC gene cassette in the E. coli genome

First, the TatA^eGFP^BC gene cassette was amplified by PCR using forward primer (5’-GCTCGGCTCCATCGGTTCCG-3’) and reverse primer (5’-CAGTACATCGGGATCGCCAACAGC-3’) from plasmid pBAD33-TatA-GFP-BC (in-house collection). Then, flanking regions of Tat operon were amplified from the chromosomal DNA of *E. coli* by PCR using an upstream forward flanking primer (5’-CGACAGTTTGCGCCAGGGCA-3’) and a reverse flanking primer (5’-AAGCCTTTGATCGACGCACCAAGA-3’) and a downstream forward flanking primer (5’-CGCCGGATGTCTTCTCGCAAAC-3’) and a reverse flanking primer (5’-AAGTCGCAGCTTGCCACTGGC-3’). The resulting flanking fragments and TatA^eGFP^BC gene cassette were incubated for 1 hour in 1.33X Gibson mastermix (Gibson, Young et al., 2009). The resulting TatA^eGFP^BC gene cassette fused with flanking regions was amplified using PCR. Finally, the amplified gene product was introduced into the *E. coli* MC4100 strain containing the chromosomal CAT/SacB cassette and the pSIM6 plasmid using electroporation, as described above. To check whether the recombination of TatA^eGFP^BC was successful, the cells were grown and plated on plates containing 5% sucrose and plates containing chloramphenicol (15 μg/ml). Cells that grew on sucrose and died on chloramphenicol were screened by PCR and sequenced to identify positive TatA^eGFP^BC colonies. We identified one completely correct strain, and one in which accidentally the first part of the *tatB* gene was deleted during recombination. The latter strain was used as a negative control in some of the subsequent experiments.

### Western-blot functionality assay

To check the transport function of fluorescently labeled TatA, the endogenous *E. coli* Tat substrate protein SufI was fused at its carboxy-terminus to a flexible Pro-Gly-Gly linker and then the influenza hemagglutinin tag (HA-tag), and cloned into plasmid pCL1920 using standard molecular biology techniques. Plasmid pCL1920-SufI-HA was then transformed into ΔTatABC, TatA^eGFP^BC, Jarv16 (MC4100 ΔtatA/E) (Sargent, Stanley et al., 1999) and wild-type MC4100 *E. coli* strains. Transformants were selectively grown on LB plates containing Spectinomycin (50 μg/ml). All four Tat variant strains were grown to mid-log phase at 30°C and expression of SufI-HA substrate protein was induced with 1mM Isopropyl β-D-1-thiogalactopyranoside (IPTG) for 2h at 37°C. Cells were harvested by centrifugation (6,500xg, 5 min) and the pellet was resuspended in phosphate-buffered saline pH 7.4 (PBS) to a final OD_600_ of 10. The cell suspension was diluted twice in 2X SDS sample buffer and incubated for 10 minutes in boiling water. Protein samples were separated on a 10% SDS-PAGE gel and blotted on a nitrocellulose membrane for western blot analysis. Blots were incubated with blocking buffer (PBS with 0.5% skimmed milk and 0.1% (v/v) Tween 20) for at least 4h. Primary antibody (rabbit anti-HA, Sigma H6908) was diluted 1:5000 in blocking buffer and incubated overnight. After washing, the blots were incubated for 1 hour with secondary antibody (goat anti-rabbit IgG-HRP, Promega W4018) that was diluted 1:10000 in blocking buffer. The binding of antibodies to the blot was visualized using Pierce ECL Western Blotting Substrate (Thermo Scientific). The indicated relative molecular weight of the protein was deduced from Precision Plus Protein Standards (BioRad).

### Expression of Tat substrate protein SufI and HiPIP-R10 in TatA^eGFP^BC E. coli strain

To study the effect of substrate overexpression on TatA dynamics, Tat transportable substrate protein SufI was expressed from plasmid pBAD33-SufI and the Tat translocation arresting substrate protein HiPIP-R10 - with a long flexible linker between the signal peptide and the mature domain - was expressed from plasmid pBW-R10 (Richter, Lindenstrauss et al., 2007). pBAD33-SufI and pBW-R10 plasmids were transformed separately into *E. coli* strain MC4100-TatA^eGFP^BC. Cells transformed with pBAD33-SufI were grown on LB plates containing chloramphenicol (34 μg/ml) and cells transformed with pBW-R10 plasmid were grown on LB plates containing ampicillin (100 μg/ml). The transformants were then inoculated in YT medium containing appropriate antibiotics and grown overnight at 220 rpm at 37°C. From the overnight culture 50 μl was added to 4950 μl fresh YT medium. Cells containing pBAD33 were grown for 30 minutes and expression of SufI was subsequently induced with 0.002%, 0.02% and 0.2% L-arabinose (w/v) for 1h at 37°C. Similarly, cells containing plasmid pBWR10 were grown for 30 minutes after which HiPIP-R10 expression was induced with 0.2% L-rhamnose (w/v) for 1h at 37°C. In both cases cells reached mid-log growth phase at the end of induction.

### Sample preparation for microscopy

*E. coli* strains (TatA^eGFP^BC wild-type, TatA^eGFP^BC overexpressing SufI and TatA^eGFP^BC overexpressing HiPIP-R10) that were grown up to mid-log phase as described above were collected by centrifugation for 2 minutes and suspended in minimal medium M9 (0.6% (w/v) Na_2_HPO_4_.2H_2_O, 0.3% (w/v) KH2PO4, 0.1% (w/v) NH_4_Cl, 0.1 % (w/v) NaCl, 0.002 M MgSO_4_, 0.4% (w/v) glucose, 0.0001 M CaCl_2_). For imaging, resuspended cells in minimal medium were immobilized on a thin agarose pad (1.5% (w/v) agarose in M9 medium) between a microscope slide and a cover slip (both plasma-cleaned). Finally, the sample chambers were sealed with VALAP (10 g Paraffin, 10 g Lanolin, 10 g Vaseline). Prior to imaging, samples were incubated on the microscope for 30 minutes to allow the cells to adjust to the imaging temperature (23 ± 1°C). Bacteria were then imaged at a constant temperature of 23°C using a custom-built epi-illuminated wide-field fluorescence microscope built around an inverted microscope body (Nikon, Eclipse Ti), equipped with an apochromatic 100x 1.49 NA TIRF oil-immersion objective and a stage top incubator system (Tokai Hit, Japan). Excitation light (wavelength 491 nm and 561 nm, intensity ~200 W/cm^2^ in the image plane) was provided by Cobolt diode-pumped solid-state lasers. Fluorescence images were taken continuously with an EMCCD camera (Andor iXon3 type 897), with an integration time of 32 ms per image, unless indicated otherwise. The total magnification was 200x, corresponding to 80 nm by 80 nm per pixel. Each strain (TatA^eGFP^BC wild-type, TatA^eGFP^BC overexpressing SufI and TatA^eGFP^BC over expressing HiPIP) was measured in three independent experiments (>80 trajectories per experiment, trajectory length ≥ 4 frames, average length ~15 time lags of 32 ms).

### Chloramphenicol treatment of TatA^eGFP^BC E. coli strain

In order to investigate the behaviour of TatA complexes in the absence of substrate proteins, TatA^eGFP^BC wild-type *E. coli* strain was grown up to mid-log phase and resuspended in minimal medium as mentioned above. Then, the resuspended cells were immobilized on a thin agarose pad containing chloramphenicol (100 μg/ml) between a microscope slide and a cover slip and the sample chambers were sealed as mentioned above. The sample was kept at room temperature for 3 hours and later imaged as described above. In the remainder of this paper, TatA^eGFP^BC wild-type strain treated with chloramphenicol is referred to as TatA^eGFP^BC-CAM.

### Fluorescence spot counting

Fluorescence images were imported into ImageJ software and the number of TatA complexes per cell was determined by counting bright fluorescent spots. For each bacterial strain (TatA^eGFP^BC wild-type, TatA^eGFP^BC overexpressing SufI, TatA^eGFP^BC overexpressing HiPIP-R10, and TatA^eGFP^BC-CAM) 100 cells were taken into account for counting. From 100 cells, average number of complexes per cell and standard errors of the mean were calculated.

### Flow cytometry

Cells were grown to early logarithmic growth phase and washed in M9 minimal medium exactly as described for fluorescence microscopy. Samples were analyzed in a BD Accuri C6 flow cytometer equipped with a 488 nm laser and a 530/30 nm emission filter, measuring forward scatter, side scatter and fluorescence intensity.

### Single-particle tracking

Data was analyzed as described in (Oswald et al., 2014). In short, images were analyzed using custom-written routines in MATLAB (MathWorks). For automated single-particle tracking in bacteria a modified version of the tracking algorithm *utrack* was used (Jaqaman, Loerke et al., 2008). In order to account for changes in the point-spread function due to axial movement of particles along the highly curved bacterial surface (in or out of focus), the location and intensity of the particles was obtained by a Gaussian fit with variable width. In addition, background subtraction was performed using a local approach allowing multi-particle localization (Leake et al., 2008). Subsequently, using the *utrack* linking algorithm, particle localizations were linked and singleparticle trajectories were reconstructed. Note that the trajectories obtained in this way are 2Dprojections of the original 3D trajectories along the highly curved *E. coli* cytoplasmic membrane. To correct for this, we used IPODD (Inverse Projection Of Displacement Distributions) for converting the 2D-projection back into the most likely 3D-displacement distribution (Oswald et al., 2014). Then, from the 3D-corrected distributions of displacement, diffusion coefficients were extracted using mean squared displacement (MSD) or cumulative probability distribution (CPD) analysis.

### MSD Analysis

For each TatA trajectory consisting of N time points, the displacement distribution *r_i,nΔt_* at a time interval *τ = nΔt* was determined using:

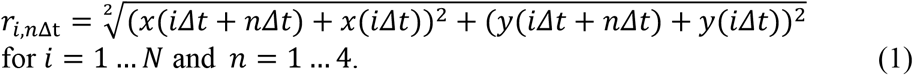

Values for *r_i,nΔt_* from all detected single-particle trajectories were pooled into a discrete 2D displacement probability distribution *PD_2D_(mΔr,τ)* for time intervals *τ ≤ 4Δt* and bin sizes ranging from 0 to 1000 nm, in *Δr* = 5 nm increments. Inverse projection of the displacement distribution (IPODD) was performed by multiplying the *PD_2D_(mΔr,τ)* with the appropriate inverted projection matrix, yielding the most probable 3D displacement probability distribution PD_3D_(τ) (Oswald et al., 2014). The average diffusion constant was determined by means of mean squared displacement (*MSD*) analysis (Martin, Forstner et al., 2002) including the experimental localization accuracy:

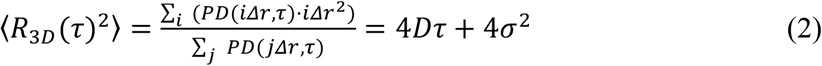

### CPD Analysis

Heterogeneity in diffusion was probed by analyzing the cumulative probability distribution (CPD). To this end, the discrete 3D-corrected displacement distribution PD (τ) for a given time-lag was integrated:

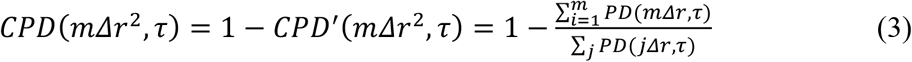

Assuming two populations simultaneously exhibiting Brownian motion the corresponding cumulative probability function (*CPF*) is expected to be the sum of two exponentials (Deverall, Gindl et al., 2005).

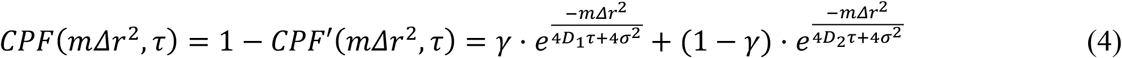

### Cell-membrane fractionation and sodium carbonate extraction of TatA-eGFP protein

Over-night precultures of *E. coli* TatA^eGFP^BC and of *E. coli* MC4100 transformed with pBad24-MscS-sfGFP (Oswald et al., 2016) were used to inoculate 500 ml of pre-warmed LB medium in a baffled Erlenmeyer flask to an OD_600_ of 0.05. The cultures were then grown to mid-log (OD_600_ of 0.5-0.6) at 30°C for improved GFP folding. The MscS^sfGFP^ expression was induced by addition of 0.1% arabinose (w/v) to the medium during the last hour of growth. The cells were pelleted for 20 minutes at 10.000 RPM (Beckman Coulter Avanti J-30I using the JA-10 rotor). The supernatants were decanted, the pellets were resuspended in 10 ml phosphate-buffered saline pH 7.4 (PBS). The cells were pelleted again for 20 minutes at 4000 RPM (Eppendorf 5810R swing out) and resuspended in 5 ml PBS containing protease inhibitors (cOmplete, pefabloc and 10 mM EDTA). Then the *E. coli* cells were lysed using a One shot cell disruptor at 1.8 kBar pressure. The cell lysates were cleared by centrifugation for 20 minutes at 10.000 g. Supernatants were then centrifuged at 278.700g for 40 minutes using a Beckman Coulter Optima Max-TL ultracentrifuge using the TLA120.2 rotor. The supernatants were collected and stored at 4°C (containing the cyto- and periplasmic fraction). The supernatant close to the membrane fraction was discarded to avoid mixing of fractions. The pellets were resuspended in either 1.2ml PBS or 1.2ml carbonate buffer (0.1 M Na2HCO3 pH 11.5), both containing protease inhibitors. The membrane pellets were homogenized using glass tissue grinders for Eppendorf tubes. The samples were incubated for 1 hour in the cold room, mildly shaking at 800 RPM (MS1 minishaker IKA). The samples were centrifuged again for 40 minutes at 278.700 g; the pellets contained the membrane fractions and the supernatants contained the washed-off proteins. Pellets were dissolved in 1.2ml PBS using glass tissue grinders for Eppendorf tubes.

### Fluorescence spectroscopy and western blot analysis

To check whether the membrane fraction and supernatant isolated using Na_2_CO_3_ and PBS contained TatA-eGFP protein, we recorded GFP fluorescence spectra using a Cary Eclipse spectrophotometer (Agilent), with excitation at 480 nm and emission recorded between 490 and 600 nm.

Samples were then mixed with 5x SDS sample buffer containing 10% freshly added β-mercaptoethanol. These were then incubated for 1 hour or overnight at 30°C. Purified enhanced Green Fluorescent Protein (His Tag) was used as positive control for the Western-blot. Protein samples were separated on 12% SDS-PAGE gel and blotted on a nitrocellulose membrane in the cold room for western blot analysis. Blots were incubated with blocking buffer (TBS with 5% skimmed milk and 0.05% (v/v) Tween 20) for at least 4h. Primary antibodies (rabbit anti-GFP, Sigma AB3080P) were diluted 1:2000 in blocking buffer and incubated for an hour at room temperature. After washing, the blots were incubated for 1h with secondary antibodies (mouse antirabbit IgG-HRP, Promega W4018) that was diluted 1:10000 in blocking buffer. The binding of antibodies to the blot was visualized using Pierce ECL Western Blotting Substrate (Thermo Scientific). The indicated relative molecular weight of the protein was deduced from Precision Plus Protein Standards (BioRad).

### Mass spectrometry

The membrane wash experiment was carried out on *E. coli* MC4100 wild-type and TatA-eGFP cells, as described above. Directly after collecting the carbonate wash fractions the pH was reduced from 11.5 to around 8.0 using 200μl of 0.5M Tris buffer with a pH of 6.8. The two samples (TatA-GFP and WT) were incubated overnight at 4°C while gently shaking at 800 rpm with 20μl anti-GFP coated agarose beads or with 5μl anti-TatA antibody (Yahr & Wickner, 2001), respectively. Protein A beads were prepared to capture anti-TatA antibody as described by the manufacturer. In short, the beads were swollen in PBS buffer, blocked with BSA and washed again in PBS. Around 70-100μl bead suspension was added to the wild-type samples incubated with anti-TatA antibody. The beads were incubated shaking at 800 rpm at 4°C for 4 hours. Both anti-GFP and anti-TatA beads were then spun down at 3000g for 5 min at 4 °C. To elute the bound antigen off the antibodies, the beads were incubated with 75μl 2% SDS and 1μl 50mM Tris pH 7.5 at 55 °C for one hour shaking at 800 rpm. The beads were spun down for 20 min at 3000g at room temperature. The supernatant was mixed with 200μl of 8 M urea in 15 mM Tris pH 8.8. The solution was transferred to a Milipore Microcon filter and spun down at 14000g for 12 min at room temperature. This washing with a denaturing buffer was repeated three more times. Subsequently, the sample was washed four times with 200μl 50mM ammonium carbonate buffer, after which it was incubated overnight with 0.7g Trypsin in 100μl ammonium carbonate buffer at 37°C while shaking at 400 rpm. The next day 100μl of 0.1% acetic acid was added to the filter, and the peptides were spun into a new clean Eppendorf tube. The liquid was evaporated using the Speed-vac for several hours and the peptides were stored in −20 °C until nano-LC MS/MS analysis. Peptides were analysed by nano-LC MS/MS using an Ultimate 3000 LC system (Dionex, Thermo Scientific) coupled to the triple TOF 5600 mass spectrometer (Sciex). The samples were dissolved in 0.1% acetic acid before measurement.

The data of the Mass spectrometry were analyzed using MaxQuant_1.5.6.5 software. The UniProt proteome database of *Escherichia coli* MG1655 was used to identify proteins from the found peptide sequences, and the eGFP sequence was added manually behind TatA.

## Acknowledgements

We thank Dr. Peter van Ulsen (Vrije Universiteit Amsterdam), Dr. Patrick Stolle and Prof. Thomas Brüser (Leibniz Universität Hannover) for plasmids, Timothy Yahr (University of Iowa) for the anti-TatA antibody, Thomas Flor for generating ΔTatABC and TatA^eGFP^BC strains, Valentina Aiosa for imaging and Anna Chertkova, Emile Zwering and Päivi Reckman for performing and analyzing sodium carbonate extraction of membrane proteins.

We acknowledge financial support from LaserLaB Amsterdam, the Netherlands Organisation for Scientific Research (NWO) with a Vici grant and an NWO-Groot grant, and a Dutch Technology Foundation (STW) research program “Nanoscopy” grant.

The authors declare no conflict of interests.

## Supporting information

**Supplementary Figure 1.**
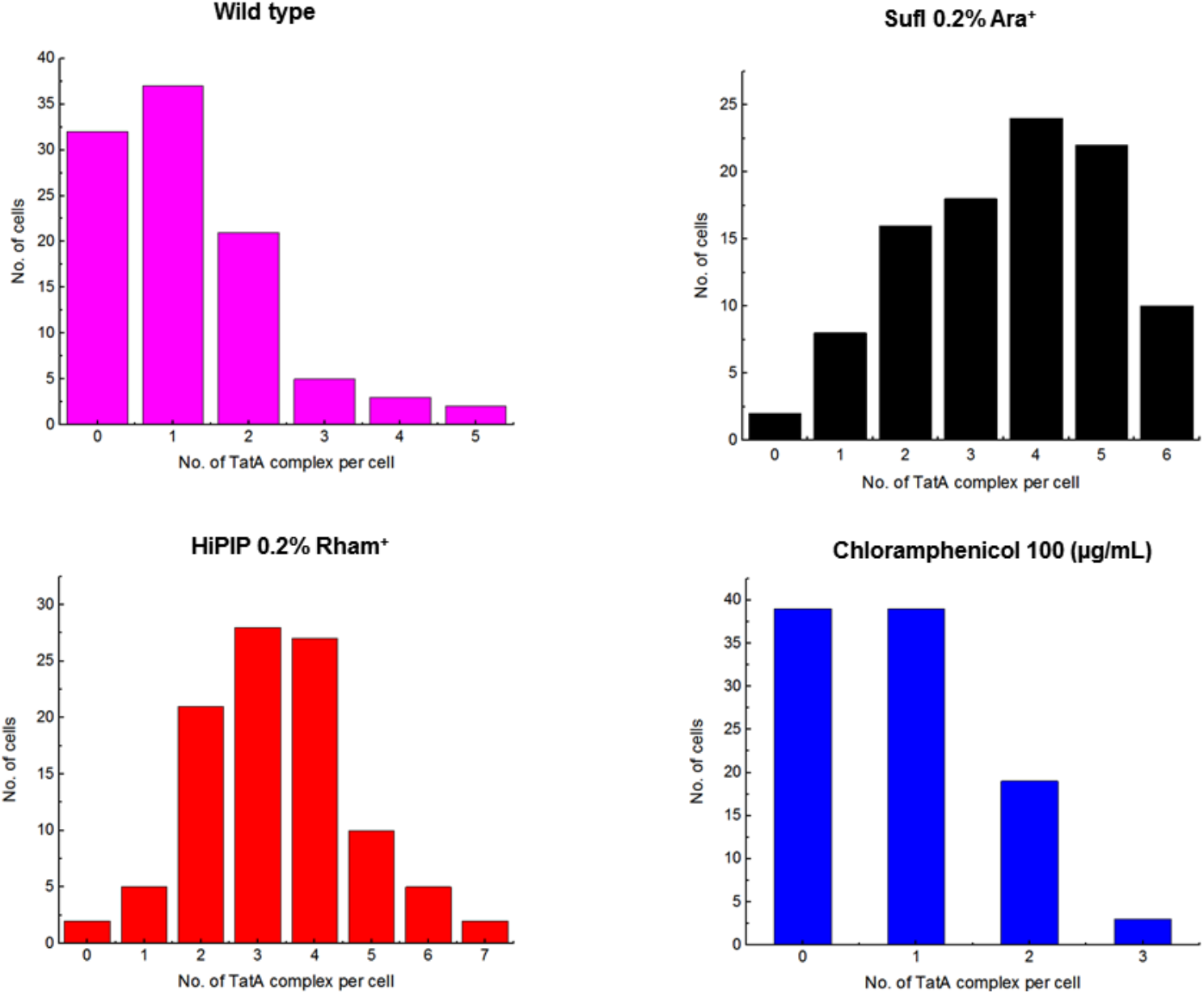
Number of TatA-eGFP complex formed in MC4100 *E. Coli* cells. TatA^eGFP^BC *E. coli* strain expressing wild-type levels of substrate (top left); overexpressing the Tat substrate SufI (top right); overexpressing the transport-arresting substrate HiPIP-R10 (bottom left); treated with chloramphenicol (bottom right).

**Supplementary Figure 2.**
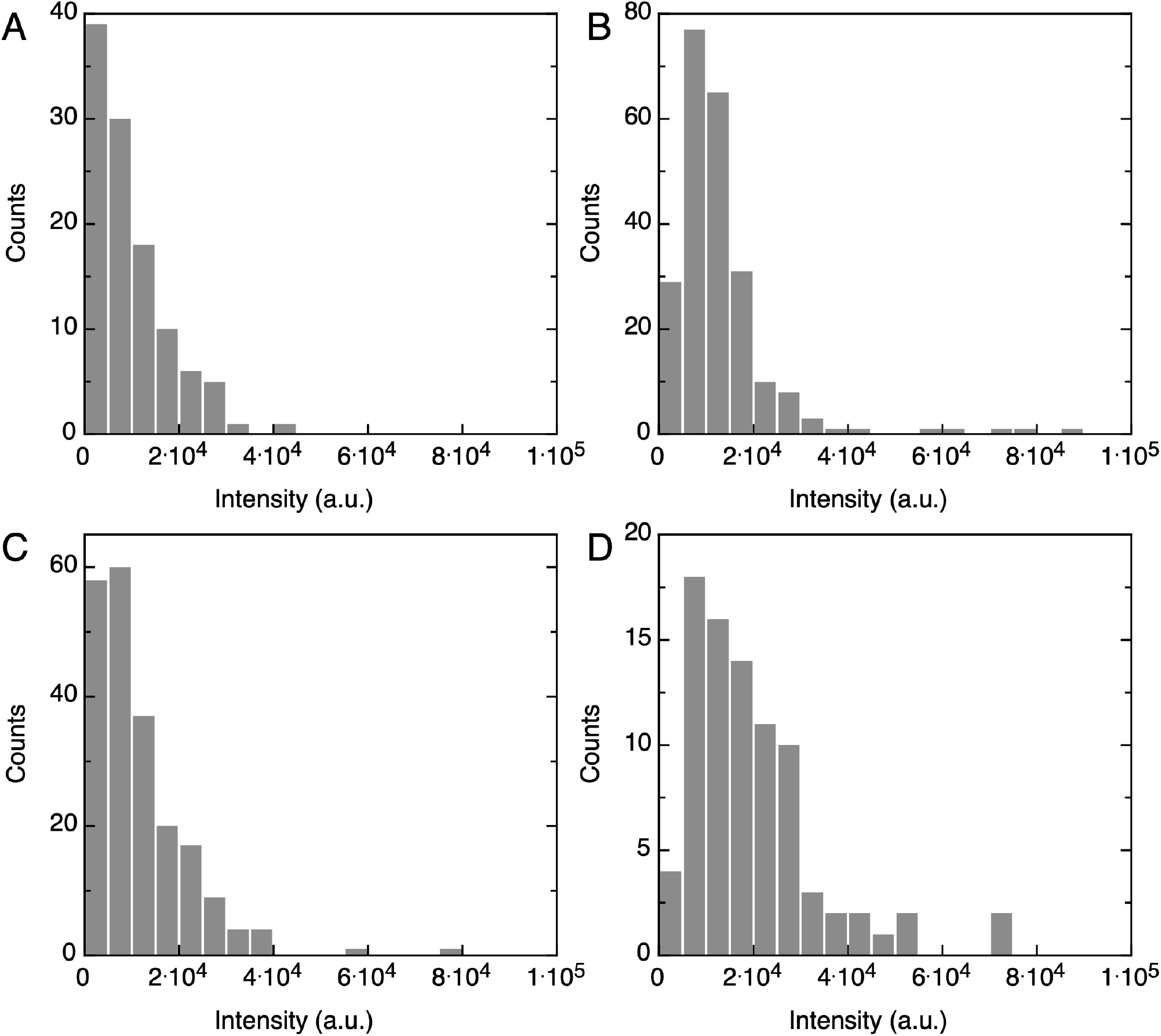
Histogram of the fluorescence intensities of tracked TatA-eGFP complexes *in vivo*. TatA^eGFP^BC *E. coli* strain expressing wild-type levels of substrate (A); overexpressing the Tat substrate SufI induced with 0.2 % arabinose (B); overexpressing the transport-arresting substrate HiPIP-R10 induced with 0.2% rhamnose (C); treated with 100 μg/ml chloramphenicol (D). For comparison, the mean intensity of purified eGFP under identical imaging conditions is 510 ± 150 a.u. (*N*=183).

**Supplementary Figure 3.**
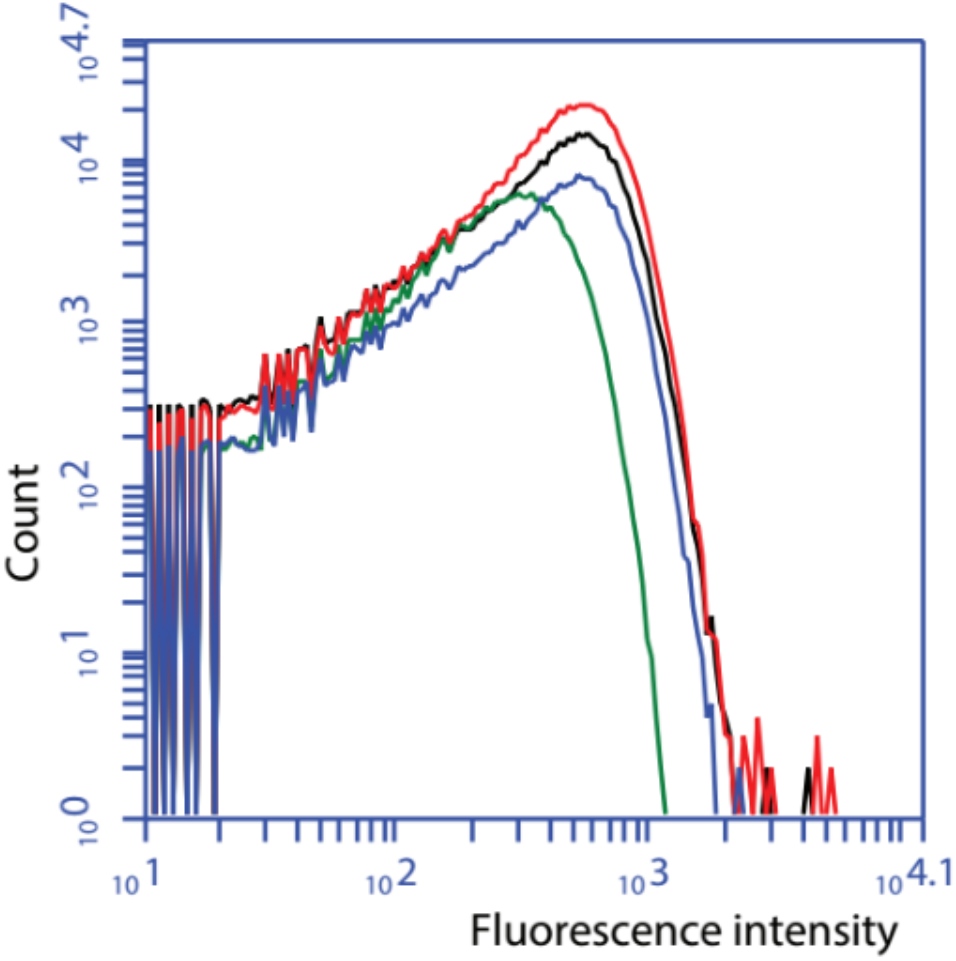
Total cellular fluorescence intensity measured using a flow cytometer. Wild-type *E. coli* MC4100 strain (green); TatA^eGFP^BC *E. coli* strain expressing wild-type levels of substrate (red); TatA^eGFP^BC *E. coli* strain overexpressing the Tat substrate SufI (black); TatA^eGFP^BC *E. coli* strain overexpressing the transport-arresting substrate HiPIP-R10 (blue).

**Supplementary Figure 4.**
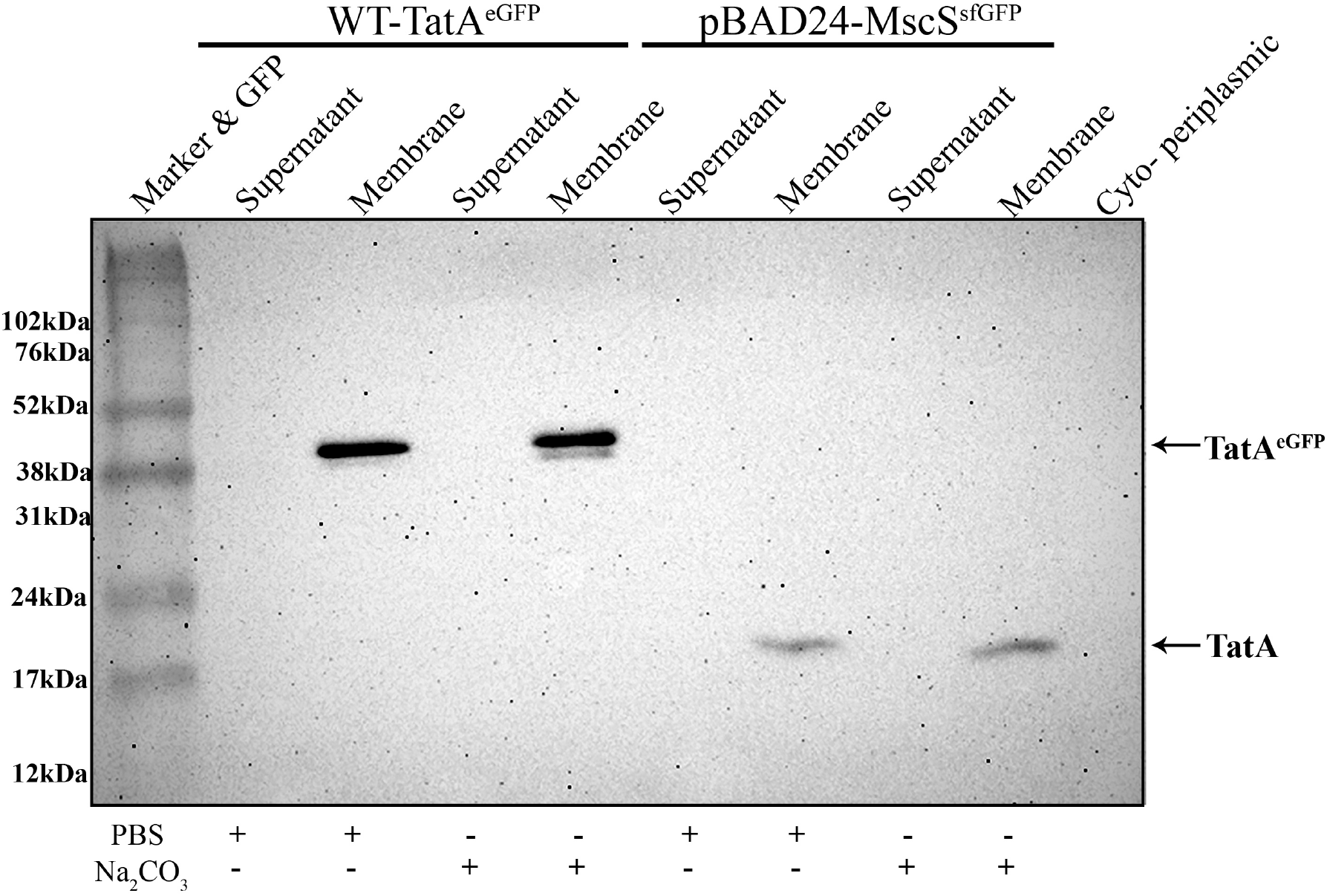
The anti-TatA antibody does not recognize TatA(-eGFP) that is removed from the membrane by carbonate extraction. Samples from Figure 4 are now probed with an anitbody against TatA. TatA is only detected in the membranes, and not in the supernatant samples in which TatA-eGFP is detected by the antibody against GFP (Figure 4).

**Supplementary Figure 5.**
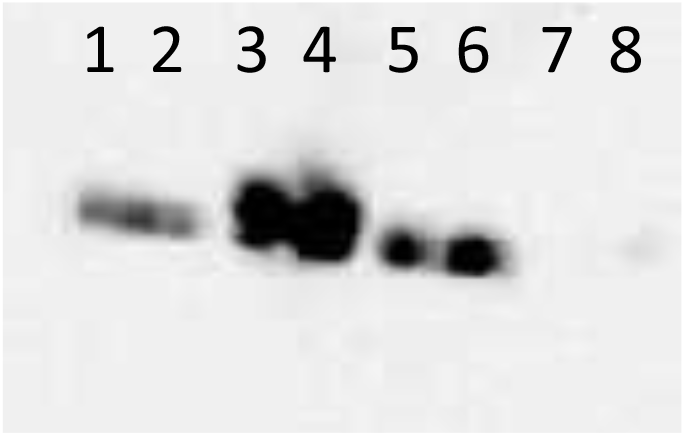
TatA-mCherry can be extracted from the membrane by carbonate wash. Western blot against mCherry using the Abcam Rabbit polyclonal antibody against mCherry (ab167453). Membranes are isolated and washed as described for TatA-eGFP. Experiment performed in duplicate. 1, 2: pellet fraction of carbonate wash; 3,4: pellet fraction of PBS wash; 5, 6: supernatant fraction of carbonate wash; 7, 8: supernatant fraction of PBS wash.

**Supplementary movie 1.**
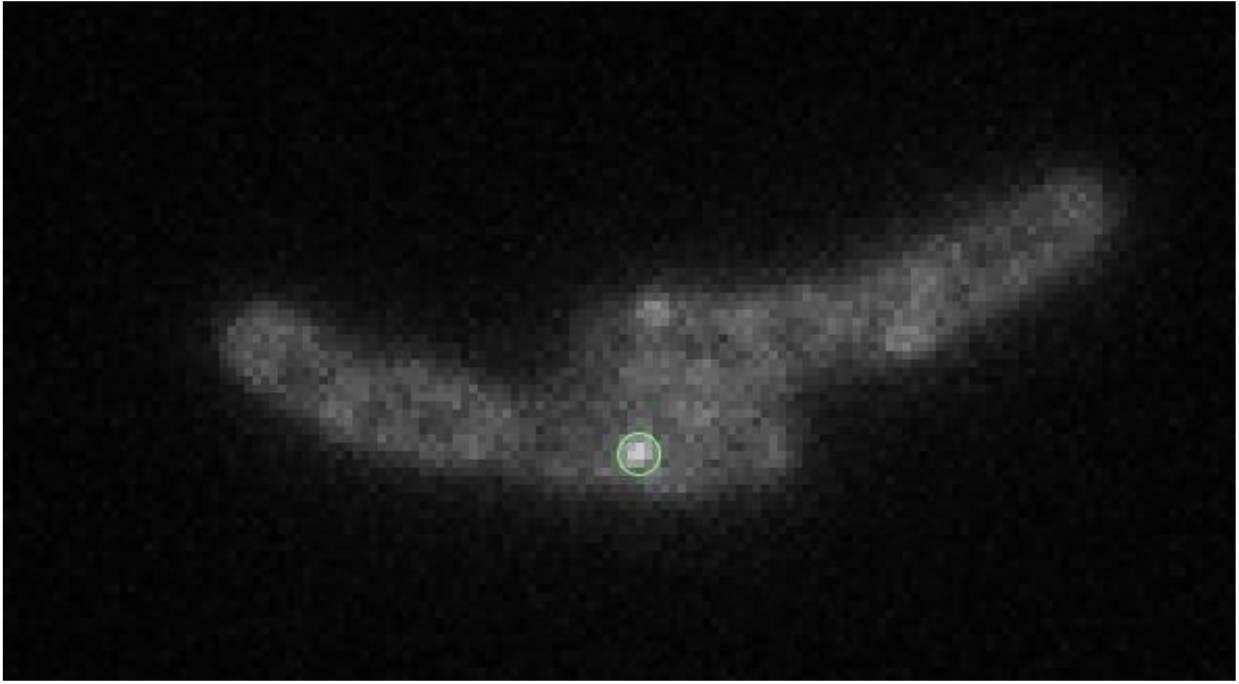
Example of a fluorescence movie of two cells expressing TatA-eGFP. The movie is recorded with 32 ms exposure time. Fluorescent spots are tracked. Movie genrated using ImageJ (Schneider, Rasband et al., 2012) and the tracking plugin Trackmate (Tinevez, Perry et al., 2017).

**Supplementary movie 2.**
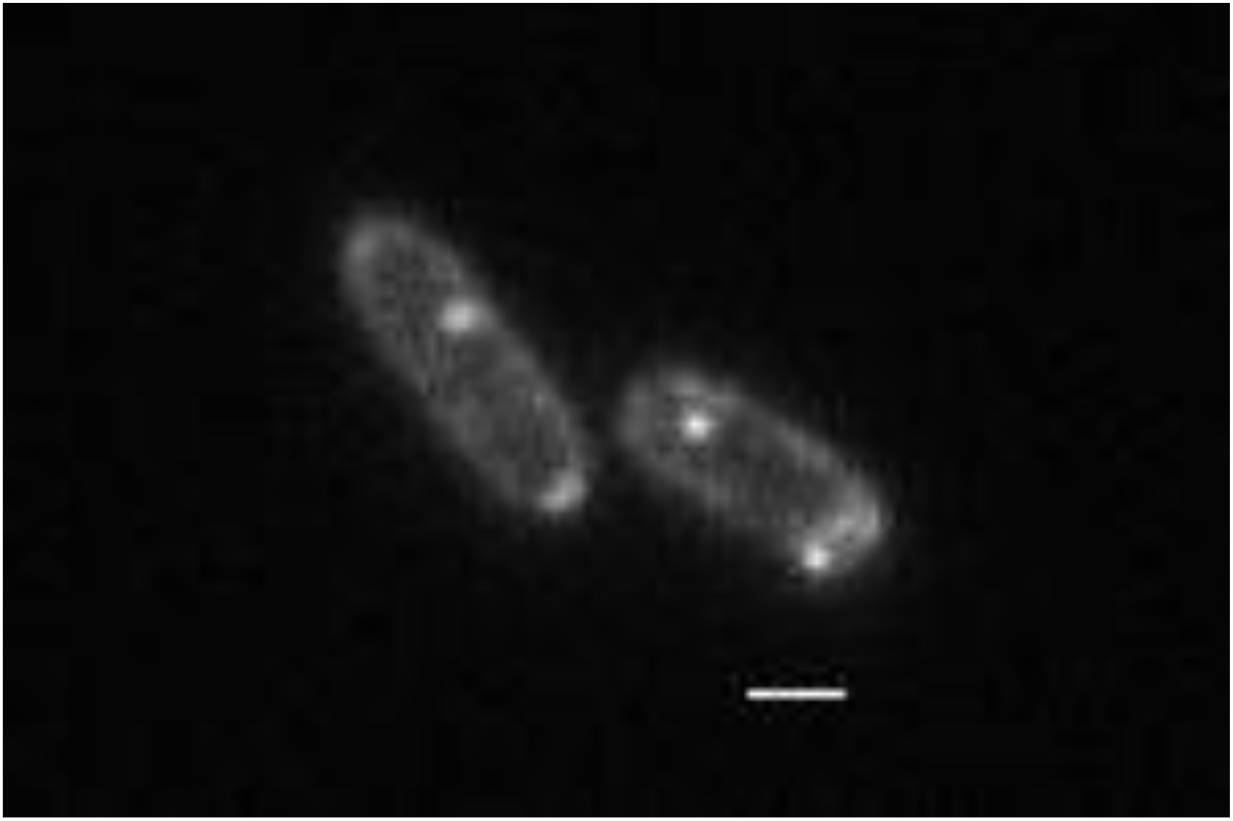
Example of a fluorescence movie of a cell expressing TatA-eGFP exposed to Chloramphenicol (100 μg/ml) for one hour. Chloramphenicol inhibits the synthesis of new substrate protein molecules. Exposure time was 32 ms. The scale bar in the first frame corresponds to 1 μm. Movie genrated using ImageJ (Schneider et al., 2012).

## References

Alami M, Luke I, Deitermann S, Eisner G, Koch HG, Brunner J, Muller M (2003) Differential interactions between a twin-arginine signal peptide and its translocase in Escherichia coli. Molecular cell 12: 937–46

Alcock F, Baker MA, Greene NP, Palmer T, Wallace MI, Berks BC (2013) Live cell imaging shows reversible assembly of the TatA component of the twin-arginine protein transport system. Proceedings of the National Academy of Sciences of the United States of America 110: E3650–9

Alcock F, Stansfeld PJ, Basit H, Habersetzer J, Baker MA, Palmer T, Wallace MI, Berks BC (2016) Assembling the Tat protein translocase. Elife 5

Bolhuis A, Mathers JE, Thomas JD, Barrett CM, Robinson C (2001) TatB and TatC form a functional and structural unit of the twin-arginine translocase from Escherichia coli. The Journal of biological chemistry 276: 20213–9

Brüser T, Sanders C (2003) An alternative model of the twin arginine translocation system. Microbiol Res 158: 7–17

Cline K, Mori H (2001) Thylakoid DeltapH-dependent precursor proteins bind to a cpTatC-Hcf106 complex before Tha4-dependent transport. The Journal of cell biology 154: 719–29

Datta S, Costantino N, Court DL (2006) A set of recombineering plasmids for gramnegative bacteria. Gene 379: 109–15

De Keersmaeker S, Van Mellaert L, Lammertyn E, Vrancken K, Anne J, Geukens N (2005a) Functional analysis of TatA and TatB in Streptomyces lividans. Biochem Biophys Res Commun 335: 973–82

De Keersmaeker S, Van Mellaert L, Schaerlaekens K, Van Dessel W, Vrancken K, Lammertyn E, Anne J, Geukens N (2005b) Structural organization of the twin-arginine translocation system in Streptomyces lividans. FEBS Lett 579: 797–802

De Leeuw E, Porcelli I, Sargent F, Palmer T, Berks BC (2001) Membrane interactions and self-association of the TatA and TatB components of the twin-arginine translocation pathway. FEBS Lett 506: 143–8

Deich J, Judd EM, McAdams HH, Moerner WE (2004) Visualization of the movement of single histidine kinase molecules in live Caulobacter cells. Proc Natl Acad Sci U S A 101: 15921–6

Deverall MA, Gindl E, Sinner EK, Besir H, Ruehe J, Saxton MJ, Naumann CA (2005) Membrane lateral mobility obstructed by polymer-tethered lipids studied at the single molecule level. Biophysical journal 88: 1875–86

Eimer E, Frobel J, Blummel AS, Muller M (2015) TatE as a Regular Constituent of Bacterial Twin-arginine Protein Translocases. The Journal of biological chemistry 290: 29281–9

Frobel J, Rose P, Muller M (2012) Twin-arginine-dependent translocation of folded proteins. Philosophical transactions of the Royal Society of London Series B, Biological sciences 367: 1029–46

Gerard F, Cline K (2007) The thylakoid proton gradient promotes an advanced stage of signal peptide binding deep within the Tat pathway receptor complex. The Journal of biological chemistry 282: 5263–72

Gibson DG, Young L, Chuang RY, Venter JC, Hutchison CA, 3rd, Smith HO (2009) Enzymatic assembly of DNA molecules up to several hundred kilobases. Nature methods 6: 343–5

Gohlke U, Pullan L, McDevitt CA, Porcelli I, de Leeuw E, Palmer T, Saibil HR, Berks BC (2005) The TatA component of the twin-arginine protein transport system forms channel complexes of variable diameter. Proc Natl Acad Sci U S A 102: 10482–6

Hou B, Heidrich ES, Mehner-Breitfeld D, Brüser T (2018) The TatA component of the twin-arginine translocation system locally weakens the cytoplasmic membrane of Escherichia coli upon protein substrate binding. The Journal of biological chemistry 293: 7592–7605

Jaqaman K, Loerke D, Mettlen M, Kuwata H, Grinstein S, Schmid SL, Danuser G (2008) Robust single-particle tracking in live-cell time-lapse sequences. Nat Methods 5: 695–702

Jong WS, ten Hagen-Jongman CM, Genevaux P, Brunner J, Oudega B, Luirink J (2004) Trigger factor interacts with the signal peptide of nascent Tat substrates but does not play a critical role in Tat-mediated export. Eur J Biochem 271: 4779–87

Kim H, Botelho SC, Park K, Kim H (2015) Use of carbonate extraction in analyzing moderately hydrophobic transmembrane proteins in the mitochondrial inner membrane. Protein Sci 24: 2063–9

Kreutzberger AJB, Ji M, Aaron J, Mihaljevic L, Urban S (2019) Rhomboid distorts lipids to break the viscosity-imposed speed limit of membrane diffusion. Science 363

Leake MC, Greene NP, Godun RM, Granjon T, Buchanan G, Chen S, Berry RM, Palmer T, Berks BC (2008) Variable stoichiometry of the TatA component of the twin-arginine protein transport system observed by in vivo single-molecule imaging. Proc Natl Acad Sci U S A 105: 15376–81

Lee PA, Tullman-Ercek D, Georgiou G (2006) The bacterial twin-arginine translocation pathway. Annual review of microbiology 60: 373–95

Lindenstrauss U, Brüser T (2009) Tat transport of linker-containing proteins in Escherichia coli. FEMS microbiology letters 295: 135–40

Martin DS, Forstner MB, Kas JA (2002) Apparent subdiffusion inherent to single particle tracking. Biophysical journal 83: 2109–17

Molloy MP (2008) Isolation of bacterial cell membranes proteins using carbonate extraction. Methods in molecular biology 424: 397–401

Natale P, Brüser T, Driessen AJ (2008) Sec- and Tat-mediated protein secretion across the bacterial cytoplasmic membrane--distinct translocases and mechanisms. Biochimica et biophysica acta 1778: 1735–56

Oates J, Barrett CM, Barnett JP, Byrne KG, Bolhuis A, Robinson C (2005) The Escherichia coli twin-arginine translocation apparatus incorporates a distinct form of TatABC complex, spectrum of modular TatA complexes and minor TatAB complex. Journal of molecular biology 346: 295–305

Oswald F, Bank ELM, Bollen YJM, Peterman EJG (2014) Imaging and quantification of trans-membrane protein diffusion in living bacteria. Physical chemistry chemical physics: PCCP 16: 12625–34

Oswald F, Varadarajan A, Lill H, Peterman EJG, Bollen YJM (2016) MreB-Dependent Organization of the E. coli Cytoplasmic Membrane Controls Membrane Protein Diffusion. Biophysical journal 110: 1139–49

Palmer T, Berks BC (2012) The twin-arginine translocation (Tat) protein export pathway. Nature reviews Microbiology 10: 483–96

Pop OI, Westermann M, Volkmer-Engert R, Schulz D, Lemke C, Schreiber S, Gerlach R, Wetzker R, Muller JP (2003) Sequence-specific binding of prePhoD to soluble TatAd indicates protein-mediated targeting of the Tat export in Bacillus subtilis. The Journal of biological chemistry 278: 38428–36

Porcelli I, de Leeuw E, Wallis R, van den Brink-van der Laan E, de Kruijff B, Wallace BA, Palmer T, Berks BC (2002) Characterization and membrane assembly of the TatA component of the Escherichia coli twin-arginine protein transport system. Biochemistry 41: 13690–7

Pradel N, Ye C, Livrelli V, Xu J, Joly B, Wu LF (2003) Contribution of the twin arginine translocation system to the virulence of enterohemorrhagic Escherichia coli O157:H7. Infection and immunity 71: 4908–16

Richter S, Lindenstrauss U, Lucke C, Bayliss R, Brüser T (2007) Functional Tat transport of unstructured, small, hydrophilic proteins. The Journal of biological chemistry 282: 33257–64

Rodriguez F, Rouse SL, Tait CE, Harmer J, De Riso A, Timmel CR, Sansom MS, Berks BC, Schnell JR (2013) Structural model for the protein-translocating element of the twin-arginine transport system. Proceedings of the National Academy of Sciences of the United States of America 110: E1092–101

Rose P, Frobel J, Graumann PL, Muller M (2013) Substrate-dependent assembly of the Tat translocase as observed in live Escherichia coli cells. PloS one 8: e69488

Saffman PG, Delbruck M (1975) Brownian motion in biological membranes. Proceedings of the National Academy of Sciences of the United States of America 72: 3111–3

Sargent F, Stanley NR, Berks BC, Palmer T (1999) Sec-independent protein translocation in Escherichia coli. A distinct and pivotal role for the TatB protein. The Journal of biological chemistry 274: 36073–82

Schneider CA, Rasband WS, Eliceiri KW (2012) NIH Image to ImageJ: 25 years of image analysis. Nature methods 9: 671–5

Shanmugham A, Wong Fong Sang HW, Bollen YJM, Lill H (2006) Membrane binding of twin arginine preproteins as an early step in translocation. Biochemistry 45: 2243–9

Sharan SK, Thomason LC, Kuznetsov SG, Court DL (2009) Recombineering: a homologous recombination-based method of genetic engineering. Nature protocols 4: 206–23

Sonnleitner A, Schutz GJ, Schmidt T (1999) Free Brownian motion of individual lipid molecules in biomembranes. Biophysical journal 77: 2638–42

Stanley NR, Findlay K, Berks BC, Palmer T (2001) Escherichia coli strains blocked in Tat-dependent protein export exhibit pleiotropic defects in the cell envelope. Journal of bacteriology 183: 139–44

Stanley NR, Palmer T, Berks BC (2000) The twin arginine consensus motif of Tat signal peptides is involved in Sec-independent protein targeting in Escherichia coli. The Journal of biological chemistry 275: 11591–6

Taubert J, Brüser T (2014) Twin-arginine translocation-arresting protein regions contact TatA and TatB. Biological chemistry 395: 827–36

Taubert J, Hou B, Risselada HJ, Mehner D, Lunsdorf H, Grubmuller H, Brüser T (2015) TatBC-independent TatA/Tat substrate interactions contribute to transport efficiency. PloS one 10: e0119761

Thomason L, Court DL, Bubunenko M, Costantino N, Wilson H, Datta S, Oppenheim A (2007) Recombineering: genetic engineering in bacteria using homologous recombination. Current protocols in molecular biology / edited by Frederick M Ausubel [et al] Chapter 1: Unit 1 16

Tinevez JY, Perry N, Schindelin J, Hoopes GM, Reynolds GD, Laplantine E, Bednarek SY, Shorte SL, Eliceiri KW (2017) TrackMate: An open and extensible platform for singleparticle tracking. Methods 115: 80–90

van den Wildenberg SM, Bollen YJM, Peterman EJG (2011) How to quantify protein diffusion in the bacterial membrane. Biopolymers 95: 312–21

Xiong Y, Santini CL, Kan B, Xu J, Filloux A, Wu LF (2007) Expression level of heterologous tat genes is crucial for in vivo reconstitution of a functional Tat translocase in Escherichia coli. Biochimie 89: 676–85

Yahr TL, Wickner WT (2001) Functional reconstitution of bacterial Tat translocation in vitro. EMBO J 20: 2472–9

Yu D, Ellis HM, Lee EC, Jenkins NA, Copeland NG, Court DL (2000) An efficient recombination system for chromosome engineering in Escherichia coli. Proceedings of the National Academy of Sciences of the United States of America 97: 5978–83

